# Chromothripsis orchestrates leukemic transformation in blast phase MPN through targetable amplification of *DYRK1A*

**DOI:** 10.1101/2023.12.08.570880

**Authors:** CK Brierley, BH Yip, G Orlando, H Goyal, S Wen, J Wen, MF Levine, G M Jakobsdottir, A Rodriguez-Meira, A Adamo, M Bashton, A Hamblin, SA Clark, J O’Sullivan, L Murphy, AA Olijnik, A Cotton, S Narina, SM Pruett-Miller, A Enshaei, C Harrison, M Drummond, S Knapper, A Tefferi, I Antony-Debré, S Thongjuea, DC Wedge, S Constantinescu, E Papaemmanuil, B Psaila, JD Crispino, AJ Mead

**Author notes:** contributed equally (CK Brierley, BH Yip, G Orlando). jointly supervised this work (B Psaila, JD Crispino, AJ Mead).

## Abstract

Chromothripsis, the process of catastrophic shattering and haphazard repair of chromosomes, is a common event in cancer. Whether chromothripsis might constitute an actionable molecular event amenable to therapeutic targeting remains an open question. We describe recurrent chromothripsis of chromosome 21 in a subset of patients in blast phase of a myeloproliferative neoplasm (BP-MPN), which alongside other structural variants leads to amplification of a region of chromosome 21 in ∼25% of patients (‘chr21amp’). We report that chr21amp BP-MPN has a particularly aggressive and treatment-resistant phenotype. The chr21amp event is highly clonal and present throughout the hematopoietic hierarchy. *DYRK1A*, a serine threonine kinase and transcription factor, is the only gene in the 2.7Mb minimally amplified region which showed both increased expression and chromatin accessibility compared to non-chr21amp BP-MPN controls. We demonstrate that *DYRK1A* is a central node at the nexus of multiple cellular functions critical for BP-MPN development, including DNA repair, STAT signalling and BCL2 overexpression. *DYRK1A* is essential for BP-MPN cell proliferation *in vitro* and *in vivo*, and DYRK1A inhibition synergises with BCL2 targeting to induce BP-MPN cell apoptosis. Collectively, these findings define the chr21amp event as a prognostic biomarker in BP-MPN and link chromothripsis to a druggable target.

---

The term chromothripsis describes a massive genomic rearrangement event caused by shattering and haphazard realignment of a chromosomal region.^1^ Recent large pan-cancer studies have identified that chromothripsis is pervasive across solid tumors, occurring in 29-42% of cancers – a far higher incidence than previously appreciated – and is associated with an adverse prognosis.^2,3^ Chromothripsis is associated with defective DNA repair pathways, including a strong association with mutation of *TP53* (m*TP53*), although 60% of chromothripsis cases occur in *TP53* wild-type tumors.^2^ Whilst oncogene amplification and tumor suppressor gene loss are well-described consequences of chromothripsis^2^, the mechanism and impact on disease biology conferred by specific chromothripsis events have not been elucidated. Consequently, whether chromothripsis itself constitutes an actionable and therapeutically targetable molecular event remains an open question.

The blast phase of myeloproliferative neoplasms (BP-MPN) is associated with an aggressive, treatment refractory and typically rapidly fatal disease course. BP-MPN-derived leukemias harbor a distinct molecular, morphological and clinical profile when compared to *de novo* acute myeloid leukemia (AML).^4,5^ Conventional AML treatment approaches are ineffective and the only curative treatment option is allogeneic stem cell transplant.^6^ However, only a minority of patients are eligible for transplant, and even with this approach, outcomes are poor.^7^ There is consequently a major unmet need to identify novel treatment approaches.

The mutational landscape associated with progression to BP-MPN is well-described, with frequent presence of multiple ‘high risk’ mutations that are associated with a poor prognosis in chronic phase MPN, including *ASXL1, IDH1/2, RAS, RUNX1*, spliceosome mutations and a particularly high incidence of *TP53* pathway alterations.^5,6^ Furthermore, whilst copy number alterations (CNA) and structural variants (SV) are infrequent in chronic phase MPN, these events occur with a high frequency in BP-MPN. This includes recurrent regions of deletions of 17p or 5q, monosomy 7, trisomy 8, 12q rearrangements and gains of chr1q.^8–11^ Copy number neutral-loss of heterozygosity (CNN-LOH) events affecting JAK2 and TP53 loci on 9p and 17p respectively are also well-described.^9,10,12^ However, aside from JAK2 and *IDH1/2* mutations^13–15^, few if any of these molecular events are associated with known actionable therapeutic targets.

Due to the long latency between chronic and blast phase in the majority of patients, MPN has long been studied as an examplar tractable model of genetic evolution in cancer.^16–19^ In this study we set out to identify the prevalence and downstream consequences of chromothripsis in BP-MPN, in order to determine how these events contribute to leukemic progression. Although chromothripsis has been reported to occur in ∼7% of *de novo* AML^20^, chromothripsis has not been described in BP-MPN, and the contribution of recurrent chromosome rearrangements to transformation in MPN remains poorly delineated.

We herein identify a recurrent genome amplification event (‘chr21amp’) driven by chromothripsis and other SV as a biomarker of adverse prognosis in BP-MPN. We demonstrate that chr21amp leads to upregulation of one specific gene-dual-specificity tyrosine phosphorylation-regulated kinase 1A (*DYRK1A*) - a serine threonine kinase highly conserved across evolution, which modulates a multitude of proteins critical to cellular processes such as cell cycle and quiescence, DNA damage repair, transcription, and splicing.^21–23^ *DYRK1A* acts in a dose-dependent manner, whereby overexpression and abrogation have both been demonstrated to cause disease, in a tissue and cell type-dependent manner.^24^ We propose *DYRK1A* overexpression as a new driver of progression of MPN to blast phase, and elucidate its mechanisms of impact. *DYRK1A* amplifies the basal activation of the JAK/STAT pathway seen in MPN, drives genomic instability and downregulates apoptosis, in part by increased expression of the anti-apoptotic factor *BCL2*. We demonstrate that this pathway is druggable, and that *DYRK1A* overexpression leads to a selective vulnerability to DYRK1A inhibition, which can be synergistically amplified by co-inhibition of the downstream dependency BCL2, thereby identifying a new tractable therapeutic axis for BP-AML.

## Chromothripsis-associated chr21amp is a recurrent and adverse-prognosis genome amplification event in BP-MPN

We studied a cohort of 64 BP-MPN patients with a median clinical follow-up of 6.2 months (range 0-48) and at a median age of diagnosis of 70 years (range 29-84) (**Fig 1A**, **Extended Data Table 1**). We performed integrated copy number (CN) and mutation profiling by single nucleotide polymorphism (SNP) array karyotyping and targeted sequencing of genes commonly mutated in myeloid neoplasms. Analysis of SNP array data using MoCha ^25,26^, identified that 54/64 (84.4%) of cases had CNAs, with a total of 344 CNAs and a median of 3.5 events (range 0-23) per patient. Of these, 24 (7%) were CNN-LOH events, 103 (30%) were gains and 217 (63%) losses (**Extended Data Fig 1A & B**). The majority of recurrent events had been previously described in myeloid disease, including chr1q gain (in 10/64 cases, 16%), monosomy 7 (6/64, 9%), partial or complete loss of chr5q (17/64, 27%), and loss of 17p (10/64, 16%, **Extended Data Fig 1A**). CNN-LOH on chr9p over the *JAK2* locus arose in 6 *JAK2* mutant cases, and on chr17p in 3 *TP53* mutant cases.

**Fig. 1.**
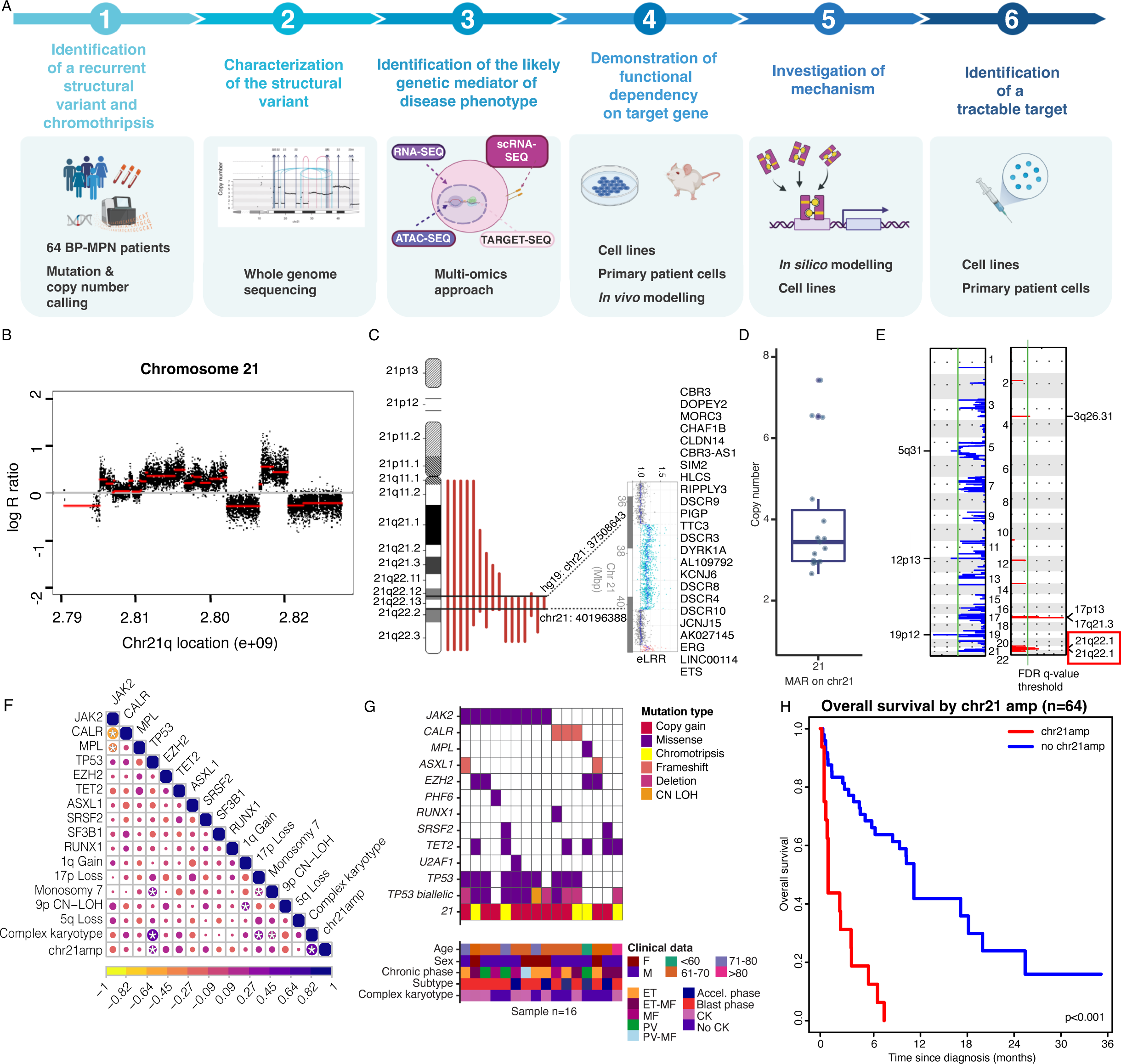
Chromothripsis-associated chr21amp is a recurrent and adverse-prognosis genome amplification event in BP-MPN. **A.** Study overview. **B.** Representative log R ratio plot of chromosome 21 derived from SNP karyotyping assay (DNACopy analysis) showing chromothripsis of chromosome 21 (’chr21amp’). **C.** Graphic displaying the minimally amplified region (‘MAR”) in common across all chr21amp cases n=16). **D.** Boxplot of median/interquartile range (IQR) of copy number overlying the chr21amp MAR for all cases (n=16, the lower and upper hinge correspond to the IQR (25th and 75th percentile), with the upper and lower whiskers extending from the hinge to +/-1.5*IQR). **E.** GISTIC analysis of recurrently lost (blue) and amplified (red) focal regions across all cases. Green horizontal line depicts the False Discovery Rate (FDR)-adjusted q-value threshold of 0.05 (n=64). **F.** Correlation plot depicting Pearson correlation coefficient of myeloid mutations and most frequent CNAs. Purple denotes positive co-variance, yellow negative, star denotes *p-val <0.05*. **G.** Mutations, TP53 allelic status, and baseline clinical data for 16 chr21amp patients. CN-LOH=Copy-neutral loss of heterozygosity, F=female, M=Male, ET=Essential Thrombocythemia, ET-MF=ET progressing to Myelofibrosis, MF=Myelofibrosis, PV=Polycythemia rubra vera, PV-MF=PV progressing to MF, Accel phase= Accelerated phase (bone marrow blast % >10 < 20%), CK=complex karyotype. **H**. Kaplan-Meier analysis of BP-MPN patients stratified by presence/absence of chr21amp event.

11/64 (17.2%) patients showed at least one chromothripsis event, a higher rate than the ∼7% incidence demonstrated in AML.^2,20,27^ Deleterious *TP53* mutations are known to be a predisposing factor for chromothripsis, with an odds ratio of around 1.54 (95% CI 1.21-1.95, p<10^-3^).^2^ As expected, there was a positive association between the presence of chromothripsis and *TP53* mutation/loss (mutation n=25,17p loss n=10, *TP53* mutation and/or loss n=29 (45.3%), chromothripsis 10/29 (34.5%), Fisher’s exact test*, p=0.002*). However, *TP53* loss was not an obligatory event for chromothripsis to occur, and 10% of cases with chromothripsis occurred in the absence of genomic alteration of p53.

We noted that a number of patients (5/64, 8%) had evidence of chromothripsis affecting chromosome 21, with focal and multiple amplifications of chr21q22-23 (**Fig 1B**). 3/5 (60%) cases harbored further chromothriptic events involving other chromosomes (chr19p, chr17p and chr22p respectively). A further 10 (totalling 16/64, 25%) had a regional CN gain event over chromosome 21q, resulting in amplification of chr21q22 (‘chr21amp’) in a quarter of patients. Overlaying of samples enabled identification of the shared minimally amplified region across all 16 cases (**Fig 1C**). This spanned 2.7Mb and containing 24 genes, with a median CN of 3.5 (range 2.7-8.3) (**Fig 1D**). On statistical testing, the amplification event affecting chr21 was significantly recurrent across the cohort (GISTIC2.0, *q-value=0.00059*, **Fig 1E**) and constituted the most common chromosome amplification event in the cohort. Patients with chr21amp had a greater number of non-chr21 CNAs compared to those without (median 6.5, range 4-15 versus median 1, range 0-16, *p-value <0.001* (*Wilcoxon rank-sum test*), **Extended Data Fig 1C**). Chr21amp occurred with a range of co-mutations and clinical phenotypes, age and sex, and significantly co-occurred with m*TP53* (**Fig 1F&G)**.

Patients with chr21amp had a particularly aggressive clinical phenotype. Overall survival was significantly impaired, with none of the patients surviving 1 year, compared to 41.8% [95% CI 28.9-60.5%] of non-chr21amp cases (*p=0.00007,* **Fig 1H**). The adverse impact of chr21amp on OS was maintained on multivariate analysis when adjusting for age, sex and high-risk molecular risk including *TP53* mutation status (and *ASXL1, EZH2, IDH1/2, SRSF2, U2AF1 Q157,* HR 4.9, p<0.001, **Extended Data Table 2** for Cox regression analysis).

Together, these data identify chr21amp as a previously unrecognised and highly prevalent copy number event occurring in BP-MPN that is associated with an adverse clinical outcome.

## Chr21amp also confers an adverse prognosis in *de novo* AML

To understand whether enrichment for chr21amp occurred more broadly in myeloid leukemias, we interrogated two published AML cohorts where copy number annotation was available, comprising 191 and 3653 *de novo* AML cases. The incidence of chr21amp was 9/191 (4.5%) in the Cancer Genome Atlas (TCGA) cohort, and 117/3653 (3.3%) in the larger UK trials cohort.^28,29^ As in our BP-MPN cohort, in the *de novo* AML context, chr21amp also co-occurred significantly with *TP53* mutations or deletions (31/117, 26.5% vs 7.7%, p<0.001, *Fisher’s exact test,* **Extended Data Fig 1D**) and complex karyotype (65/117, 55.6% vs 8.8%, p<0.001, *Fisher’s exact test),* and was associated with adverse survival in both univariable (HR 1.59 [95%CI: 1.29-1.97], p<0.001) and multivariable analysis after correction for *TP53* status (HR 1.3 [95% CI 1.1-1.7], *p=0.009,* **Extended Data Fig 1E,F**). The TCGA cohort was underpowered for a survival analysis (**Extended Data Fig 1G**). These data confirm that chr21amp is less common in *de novo* AML than BP-MPN (3- 5% vs. 25%), but where it occurs, it correlates with an adverse prognosis.

## Whole genome sequencing of chromothripsis-associated chr21amp events at high resolution

To determine the precise genetic architecture of the structural variant events that led to chr21 amplification, and confirm that this is driven by *bona fide* chromothriptic events in some cases using current criteria, we performed high-depth whole genome sequencing (WGS) in five chr21amp cases, to a median coverage of 81x [range 77-86] and purity 79% [range 58-88%] (Methods).^30^ These cases were selected on the basis of sample availability and to highlight the range of genetic events whereby chr21amp occurs, with three showing high amplifications and evidence of chromothripsis on SNP karyotyping (**Fig 2A-F**), and two comprising simpler chr21 gain events (**Extended Data Fig 2A-D**).

**Fig. 2.**
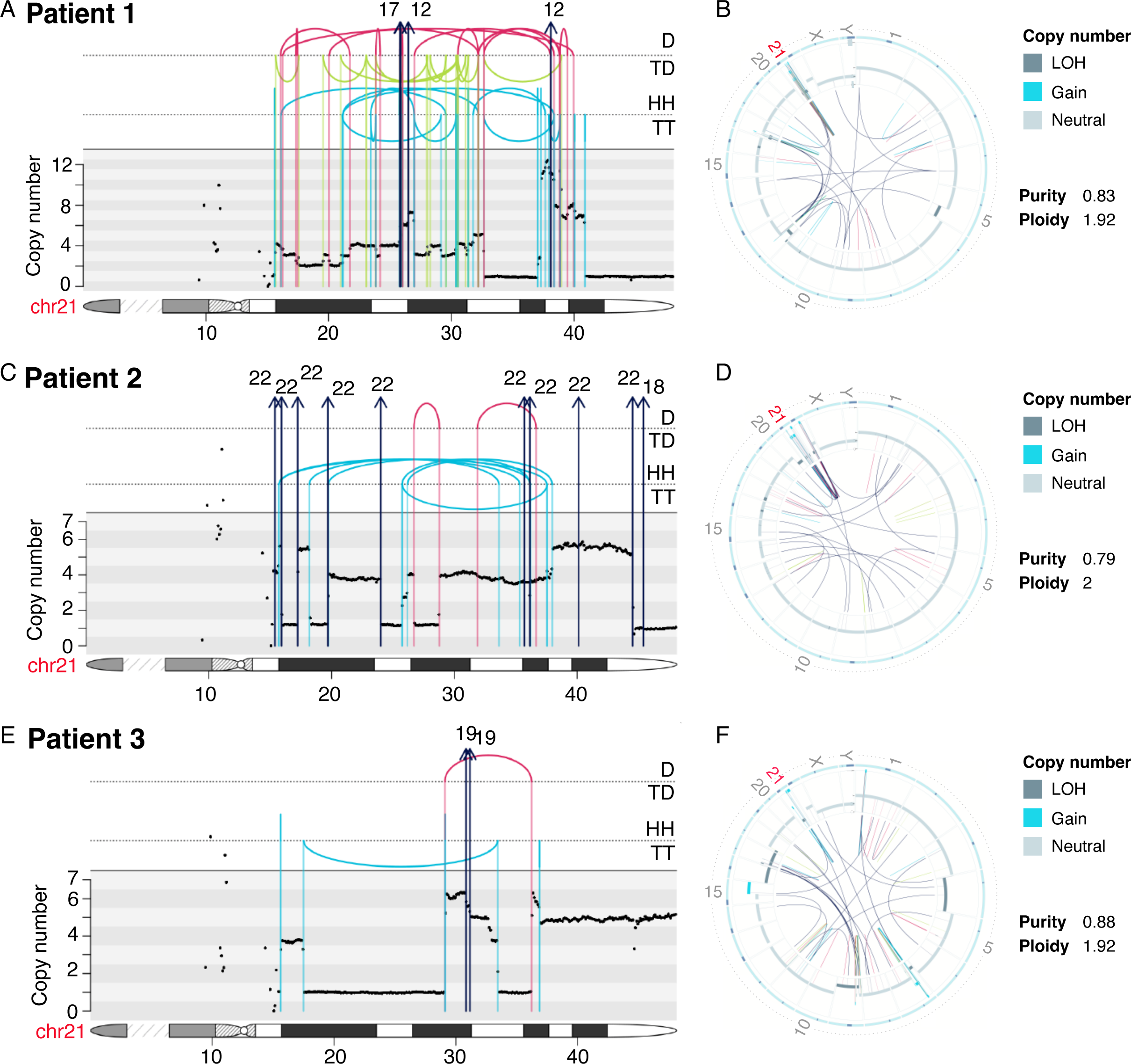
Whole genome sequencing of chromothripsis-associated chr21amp events at high resolution. **A., C. & E.** Integrated copy number and structural variant (SV) plots showing the complex SV in three chr21amp cases. The top panel shows intra-chromosomal events as arcs between breakpoint loci, and color denotes the type of SV (black=translocation, red=deletion, blue=duplication, green=inversion). Rearrangements are further separated and annotated based on orientation. D: Deletion, TD: Tandem duplication; HH, head-to-head inverted, TT: tail-to-tail inverted. Inter-chromosomal events are shown with arrows denoting the likely partner chromosome. The middle panel shows the consensus copy number across the chr21 ideogram, depicted in the lowest section of each plot to indicate breakpoint location. **B., D. & E.** Circos plots showing global SV burden corresponding to the patients in **A.,C. &E.,** demonstrating clustering around chr21. The outer ring shows the chromosome ideogram. The middle ring shows the B allelic frequency and the inner ring shows the intra-and inter-chromosomal SVs with the same color scheme as in **A.,C. & E.**

WGS provided high-resolution analysis of copy number and structural variation over the chromosome 21 amplification event, confirming and extending the SNP array findings to show that the non-recurrent translocation partners differed, with chr19 involved in 2 cases (**Fig 2E, Extended Data Fig 2C**) and chr7 (**Extended Data Fig 2A**), chr22 (**Fig 2B**) and chr17 and 12 (**Fig 2A**) implicated for others.

The genetic architecture of the structural event varied, ranging from a simple tandem duplication event (**Extended Data Fig 2C**), to multiple gains and losses along the body of chr21 (**Fig 2C**), to a highly complex amplicon involving multiple chromosomes (**Fig 2A**). Each case therefore showed a unique pattern of rearrangement. For all, chr21 formed a focus of rearrangement across the genome, as demonstrated in the circos plots (**Fig 2B,D,F, Extended Data Fig 2B & 2D**). The copy number over the shared amplified region in chr21 also varied, with a median CN of 6.5 (range 3.4-8.2). In all cases, the aberrant amplification event occurred on one allele only. Of the median 150 coding small nucleotide variants called (range 130-160), none were recurrent, and none affected the amplified region on chr21.

In the three chromothriptic cases profiled, there was a high focal density of genomic rearrangements, where numerous genomic segments were amplified distinctly and interrupted by non-amplified regions, with oscillating copy number, features in keeping with a chromothriptic events. One of the cases (**Fig 2A**) demonstrated copy number oscillations between one low (CN=2) and one very high (CN ≥ 10) event, consistent with the presence of a double minute as defined in the pan-cancer analysis of whole genomes (PCAWG) cohort.^31^ In all cases, the area of highest amplification overlay chr21q21-22.

Together, these analyses confirmed that multiple different genomic events of variable complexity converge to cause the same genetic aberration, namely copy number amplification of a specific genomic region on chr21 in the absence of an underlying mutation or recurrent chromosomal translocation. We propose that the likely explanation is amplification of a specific gene in the minimally amplified region that promotes and supports leukemic transformation.

## TARGET-seq single cell analysis prioritises gene targets amplified by the chr21amp event

We next set out to identify candidate drivers of leukemic transformation that were potentially triggered by the chr21amp event.

To enable timing of the chr21amp, clonal hierarchies and relationship to m*TP53*, we leveraged a dataset of 4 chr21amp, *JAK2* and *TP53* mutant BP-MPN patients, who had undergone TARGET-seq analysis, a multi-omic approach enabling genotype-informed analysis of copy number status and transcriptome in single cells.^19^ In brief, individual hematopoietic stem and progenitor cells (HSPCs) were isolated by index flow cytometry, enriching for early stem cell populations (Lin-CD34+CD38- CD45RA-CD90+) followed by integrated single cell genotyping at allelic resolution for m*TP53* and *JAK2* and single cell RNA sequencing (sc-RNA-seq), leveraging the softwares *inferCNV* and *numbat* to call CNAs in single cells.^32,33^ Focusing on the 4 chr21amp cases, genotyping information was available for 1903/2205 (86.3%) of cells, enabling inference of clonal hierarchy (**Fig 3A**). Due to its high sensitivity for mutation detection and flow cytometry enrichment, TARGET-seq enabled confident detection of rare precursor (residual wild-type) populations. 107 cells were detected that had no chr21amp, *TP53* or *JAK2* mutation, 179 single *JAK2* mutant cells and 162 *JAK2/TP53* co-mutant, non-chr21amp cells. Overall, chr21amp was highly clonal and co-occurred with m*JAK2* and m*TP53* in 1455/1761 (76.5%) of cells, supporting that the chr21amp event occurs after *JAK2* and *TP53* mutation (**Fig 3A**).

**Fig. 3.**
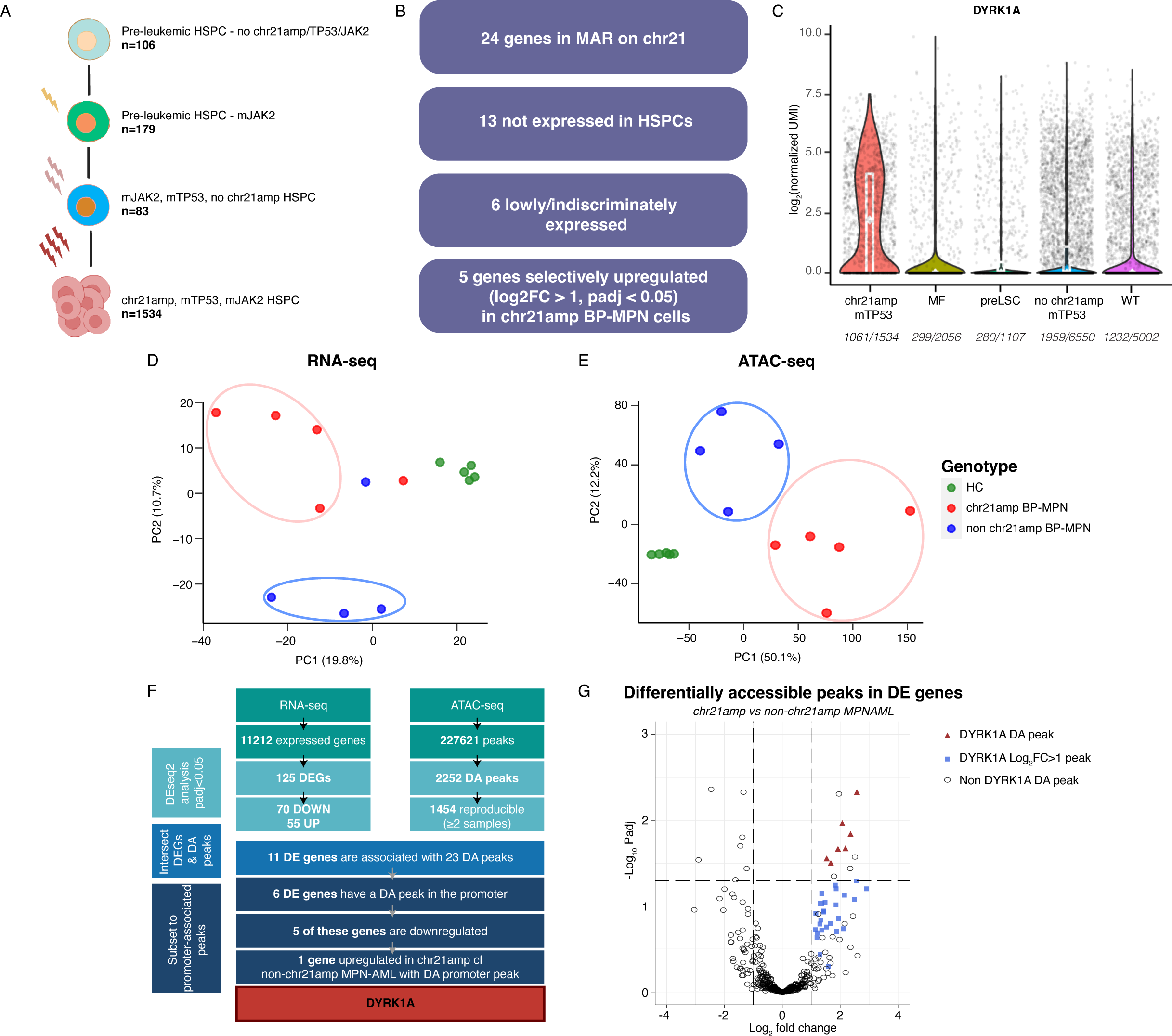
Integrated RNA- and ATAC-seq pinpoints *DYRK1A* as the putative mediator of the adverse chr21amp phenotype. **A.**TARGET-seq analysis of n=1903 cells from 4 chr21amp donors with allelic resolution of mutant JAK2/TP53 and chr21amp event in single cells enables inference of clonal hierarchy. 106 cells had no genomic aberration, while 179 cells were mutated for *JAK2V617F* alone. 83 cells were double *JAK2* and *TP53* mutant, with no evidence of chr21amp, while 1534 cells carried all 3 genomic aberrations **B.** Defining the chr21amp in single HSPCs enables prioritisation of 5 of the 24 genes in the MAR **C.** Violin plots showing that *DYRK1A* is significantly overexpressed (log2FC 2.38, p-adjusted value < 0.0001 using Benjamini Hochberg correction) in chr21amp HSPCs compared to non-chr21amp control cells including myelofibrosis (MF, n-2056 cells from 8 MF donors), pre-leukemic stem cells (preLSC, n=1107 non-mutant phenotypic HSC, identified in 12 BP-MPN donors), TP53-mutant-non-chr21amp BP-MPN (no chr21amp mTP53, n=6550 cells from 14 BP-MPN donors) and wild type cells (WT, n=5002 from 9 healthy donors). Each dot represents the expression value (log2-normalized UMI count) for a single cell, with median and quartiles shown in white. Expressing cell frequencies are shown on the bottom of each violin plot for each group. Colors are aligned with cell genotype status depicted in (**A). D.&E.** Principal component analysis of RNA-seq (**D)** and ATAC-seq (**E**) data shows clustering by chr21amp status. HC samples are depicted in green, chr21amp BP-MPN in red and non-chr21amp BP-MPN in blue. **F.** Integration of the RNA-seq and ATAC-seq datasets comparing chr21amp vs non-chr21amp BP-MPN Lin-CD34+ cells identified 125 differentially expressed genes (DEGs) and 2252 differentially accessible (DA) peaks meeting p-adjusted < 0.05. Only *DYRK1A* is DE with a DA promoter peak. **G.** Volcano plot of DA peaks in DE genes comparing chr21amp vs non-chr21amp BP-MPN samples: *DYRK1A* peaks with log2FC>1 and p-adj < 0.05 (Benjamini Hochberg correction, DESeq2 analysis) are shown in red triangles, and DYRK1A peaks with log2FC>1 are in blue squares. Of the 92 differentially accessible ATAC-seq peaks with log2FC>1, 33 occur in the *DYRK1A* gene

The clonality and late timing of chr21amp acquisition was supported by analysing the whole genome sequencing data with AmplificationTimeR.^34^ This enabled timing of the individual copy number gains occurring on chromosome 21 relative to the acquisition of somatic mutations. Within each sample, multiple gains occurred likely at the same time or in very rapid succession, findings in keeping with a single catastrophic chromothripsis event. Gains encompassing the minimally amplified region on chr21amp were universally timed as late clonal events, occurring after all mutations within the gained region, suggesting that they occurred just prior to leukemic transformation and diagnostic BP-MPN sample collection. Together, the single cell and whole genome sequencing timing analyses support that chr21amp triggers leukemic evolution.

Next, we integrated the transcriptome profiles of the chr21amp cells to prioritise genes of interest from the minimally amplified region. We compared expression in individual chr21amp HSPCs to non-chr21amp HSPCs. The non-chr21amp HSPCs included cells from normal donors, non-BP MPN donors, non-chr21amp *TP53*-mutant BP-MPN cells and non-chr21amp-non-TP53 mutant (residual wild-type/“pre-leukemic”) cells from BP-MPN donors. Of the 24 genes in this region, 13 were not expressed in HSPCs (**Fig 3B**). Four were very lowly expressed, and two (the transcription factors *ERG* and *ETS2*) were not differentially expressed across chr21amp m*TP53* BP-MPN cells and controls (**Fig 3B**). Five candidate genes (*DYRK1A, DSCR3, MORC3, PIGP, TTC3* – **Fig 3C**, **Extended Data Fig 3A-D**) were significantly upregulated and differentially expressed (log2FC>1, *padj<0.05*) in chr21amp compared to controls.

## Integrated RNA- and ATAC-seq pinpoints *DYRK1A* as the putative mediator of the adverse chr21amp phenotype

Having established that the chr21amp event was highly clonal in the HSPC compartment, to further characterize candidate genes in the amplified region we performed mini-bulk RNA-seq (n=200 cells) and ATAC-seq (n=1000 cells) on CD34+Lin-HSPC in 5 chr21amp BP-MPN patients, 4 non-chr21amp patients and 5 age-matched healthy controls. Unsupervised principal component analysis using highly variable genes and peaks in both the RNA (**Fig 3D**) and ATAC (**Fig 3E**) datasets demonstrated that healthy controls clustered tightly, and chr21amp vs non-chr21amp BP-MPN patients were stratified by principal components 1 and 2, highlighting that chr21amp accounted for a high percentage of variation and cell identity in both the gene expression and chromatin accessibility datasets.

There were 125 differentially expressed (DE) genes, of which 55 were upregulated in chr21amp vs non-chr21amp Lin-CD34+ HSPC. The only gene from the minimally amplified region that was upregulated in chr21amp cells compared to non-chr21amp BP-MPN cells was *DYRK1A* (*p value =0.0005, padj =0.03*) (**Extended Data Table 3A&B showing DESEq2 results for all 24 MAR genes on RNA and ATAC-sequencing)**. Integrated analysis of differential accessibility (DA) and DE genes comparing chr21amp *vs*. non-chr21amp Lin-CD34+ cells led to the identification of 11 differentially expressed genes associated with 23 differentially accessible peaks (**Fig 3F)**. Only *DYRK1A* was differentially expressed with a differentially accessible promoter peak (log2FC 2.36 p*adj 0.015*) - along with six further differentially accessible peaks along the gene body (*padj<0.05*) and a further 26 peaks with log2FC>1 **(Fig 3G, Extended Data Fig 3E for tracks)**.

## *DYRK1A* upregulation in other AML cohorts

Given the potential role of *DYRK1A* in driving disease transformation and adverse outcome in chr21amp BP-MPN, we sought to evaluate its role more widely in myeloid disease. Analysis of the TCGA AML cohort showed that gain of chr21q in the context of *de novo* AML is associated with increased expression of *DYRK1A* on RNA-seq (median log2FPKM 3.53 vs 3.08 [IQR 3.32-3.74 vs 2.73-3.44], *p=0.004,* **Extended Data Fig 4A**). Overexpression of *DYRK1A* in the 360-patient BEAT-AML cohort was associated with adverse OS even in the absence of chr21amp (HR 1.44, 95% CI 1.07-1.93, *p-value 0.03*, **Extended Data Fig 4B**), which was not the case for other genes in the chr21amp amplified region (**Extended Data Fig 4C&D**), implying a *DYRK1A*-specific effect. Stratifying the 360 BEAT AML patients by *DYRK1A* expression (**Extended Data Fig 4E**) into top and bottom quintiles of *DYRK1A* expression, we then performed differential gene expression analysis and GSEA. In keeping with our analyses in chr21amp BP-MPN, we observed evidence of distinct gene expression signatures associated with increased *DYRK1A* expression (**Extended Data Fig 4F**), with enrichment for multiple signaling pathways including JAK/STAT, TNFa, TGFý, and downregulation of DNA repair mechanisms (**Extended Data Fig 4G, Extended Data Table 4E&F**).

## Investigating the chr21amp associated cell state and transcriptional landscape

Next, we explored the impact of chr21amp on the transcriptional and cellular landscape in BP-MPN. Geneset enrichment analyses (GSEA) comparing DE genes in the chr21amp vs non-chr21amp Lin-CD34+ HSPC RNA-seq data showed that the top upregulated pathways were reactive oxygen species/oxidative phosphorylation along with JAK/STAT signalling, and the top downregulated pathways including those regulating cell division and survival, such as G2M checkpoint, *TP53* signaling, and apoptosis (**Fig 4A, Extended Data Table 4A&B).** GSEA between chr21amp and healthy control HSPCs demonstrated upregulation of signaling pathways including STAT5, TNFα, KRAS, PI3K and reactive oxygen species, with downregulation of G2M checkpoint and DNA repair pathways **(Fig 4B, Extended Data Table 4C&D)**.

**Fig. 4.**
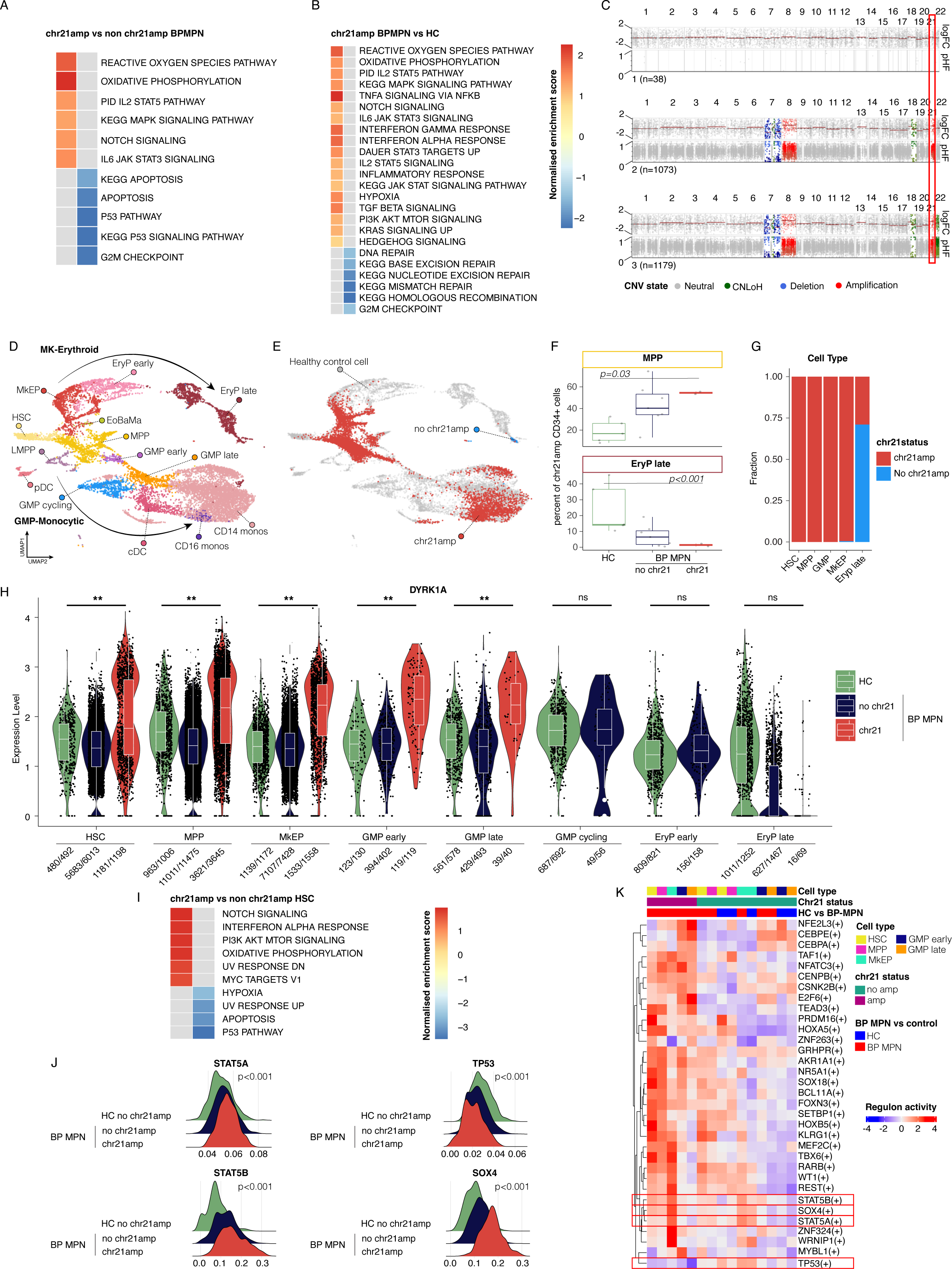
Investigating the chr21amp associated cell state and transcriptional landscape. **A.&B.** GSEA for selected KEGG and HALLMARK pathways with normalized enrichment score (NES)>1 in chr21amp (n=5) vs non-chr21amp BP-MPN (n=4) (**A.)** and chr21amp BP-MPN (n=5) vs healthy control (n=5) (**B.**) RNA-seq datasets. The heatmap shows the normalized enrichment score and the title indicates the cohort that the geneset is enriched/depleted for. Raw data can be found in **Extended Data Table 4A-D**. **C.** Clone-specific pseudobulk profile for a representative patient showing detection of the chr21amp event in single cells by the copy-number calling software *numbat*. Each of the three plot subpanels defines a clone ascertained by numbat, with the chromosomal location along the x axis, the top section showing the log2-fold change, and the bottom the paternal haplotype frequency (PHF), inferred from haplotype phasing of SNPs genotyped from single cell transcriptomes. CNV calls are colored by type of alteration (amplification in red, deletion in blue, copy-neutral loss of heterozygosity in green). The red box highlights chr21. **D.** UMAP representation of a healthy donor hematopoietic hierarchy generated by marker gene analysis of n=6134 healthy LIN- CD34+ hematopoietic stem and progenitor and myeloid cells. HSC: Haematopoietic stem cell, MkEP: Megakaryocyte-erythroid progenitors, EryP: Erythroid progenitors, EoBaMa: Eosinophil-Basophil-Mast progenitors, MPP: multipotent progenitors, LMPP: lymphoid-primed multipotent progenitors, GMP: granulocyte-monocyte progenitors, cDC: classical dendritic cell, pDC: plasmacytoid dendritic cell, monos=monocytes. **E.** UMAP projection of n=6629 cells from 2 chr21amp BP-MPN donors onto the healthy donor hematopoietic atlas colored by chr21amp status (chr21amp cell= red, non-chr21amp cell=blue, healthy control background = grey) **F.** Plots of the % of CD34+ positive cells called as MPP and EryP based on projection analysis in (e), showing expansion of MPP and depletion of EryP compared to normal controls. (**p<0.01, significance testing by paired Wilcoxon rank-sum test). **G.** Barchart depicting the fraction of cells called as chr21amp from 2 chr21amp donors, demonstrating the differentiation block into erythroid cells. **H.** Violin plots of *DYRK1A* expression in HSPC progenitors, showing transcriptional upregulation across the hematopoietic hierarchy when compared to non-chr21amp BP-MPN and HC cells. Each dot represents the expression value (log2-normalized UMI count) for each single cell, with median and quartiles shown in white. Expressing cell frequencies are shown on the bottom of each violin plot for each group. P-values calculated by Wilcoxon rank-sum test. **I.** GSEA for HALLMARK pathways with normalized enrichment score (NES)><1 in chr21amp vs non-chr21amp HSCs. The heatmap shows the normalized enrichment score and the title indicates the cohort that the geneset is enriched/depleted for. Raw data can be found in **Extended Data Table 4G&H**. **J.** SCENIC regulons scored by the AUCell algorithm upregulated in chr21amp include *STAT5A, STAT5B* and *SOX4*, with *TP53* downregulated, corroborating the GSEA findings. Statistical significance testing performed with Kruskal Wallis test with Benjamini Hochberg adjustment for multiple testing. **K.** Heatmap of SCENIC regulon analysis showing that chr21amp and *DYRK1A* overexpression leads to activation of divergent transcriptional programs. Regulons cluster by chr21amp status over cell type, when comparing chr21amp BP-MPN and non-chr21amp BP-MPN with healthy controls. STAT5A and STAT5B regulons (highlighted in red box) are upregulated in a cell-type agnostic manner by the chr21amp event, while TP53 is downregulated.

To investigate the effect of chr21amp on cell differentiation, we performed droplet-based, high-throughput single-cell RNA-seq on Lin-CD34+ cells and total mononuclear cells for 2 chr21amp patients (n=6629 cells post QC, see Methods), 9 non-chr21amp patients (n=27,492 cells) and 5 healthy bone marrow controls (n=6,143 cells). The chr21amp event was readily identified in individual cells (**Fig 4C**) and highly clonal.

Projection of the cells from chr21amp BPMPN patients onto the healthy control reference (**Fig 4D & 4E**) showed that chr21amp cells are present from the apex of the hematopoietic differentiation hierarchy, with chr21amp HSPCs particularly expanded at the multipotent progenitor (MPP)-precursor stage **(Fig 4E&F)**. Chr21amp cells were notably less frequent in late erythroid precursors, while non-chr21amp cells were exclusively detected in the late erythroid precursor cluster, implying presence of a differentiation block (**Fig 4F&G**). Along the hematopoietic differentiation hierarchy, chr21amp HSCs, MPPs, GMPs and MkEPs showed significantly elevated *DYRK1A* expression relative to non-chr21amp cells (**Fig 4H**). GSEA comparing chr21amp and *DYRK1A*-upregulated HSCs with non-chr21amp HSCs again demonstrated upregulation of multiple signaling pathways including MYC, Notch, and PI3 kinase signaling, with downregulation of apoptosis and *TP53* pathways (**Fig 4I, Extended Data Table 4G&H**).

We performed single-cell regulatory network inference and clustering (SCENIC) analysis by cell type to investigate whether the chr21amp event had a global effect on shaping active gene regulatory networks compared to non-chr21amp and healthy control HSPCs.^35,36^ This demonstrated clustering by chr21amp status, showing that chr21amp BP-MPN cells were transcriptionally distinct from both healthy control HSPCs and non-chr21amp cells (**Fig 4J&K)**. Key transcription factor networks including signalling pathways such as STAT5A and STAT5B along with negative regulators of apoptosis such as SOX4 ^37^ were globally upregulated in chr21amp HSPCs compared to both non-chr21amp BP-MPN and healthy controls (**Fig 4J**). Conversely, and in keeping with the GSEA analyses, down-regulation of the TP53 transcription factor network was observed (**Fig 4J)**.

Collectively, these findings confirmed that selective transcriptional upregulation of *DYRK1A* in chr21amp cases occurs throughout the hematopoietic hierarchy. Chr21amp was associated with increased chromatin accessibility across the *DYRK1A* locus in BP-MPN. Chr21amp and DYRK1A overexpression was associated with transcriptional evidence of perturbed cellular processes critical for transformation in MPN including cell signalling, apoptosis and DNA repair pathways. We therefore selected *DYRK1A* as the lead candidate gene for further independent validation, functional and mechanistic studies.

## *DYRK1A* expression and dependency in cell line models

We next sought to functionally validate *DYRK1A* as a gene conferring a cell survival advantage in the BP-MPN context. *In silico* screening of Broad’s Cancer Dependency Map (DepMap) shows cancer cell line dependency scores were linked to *DYRK1A* gene expression (*p<0.0001* by linear regression, **Extended Data Fig 4H**). Myeloid cell lines were amongst the highest expressors of *DYRK1A* (**Extended Data Fig 4I**) and demonstrated the highest gene dependency (**Extended Data Fig 4J**).

A kinase domain-focused CRISPR screen highlighted that both the *JAK2* mutant BP-MPN cell lines (human erytholeukemia [HEL] and the megakaryoblastic leukemia line [SET2]) are hypersensitive to *DYRK1A* targeting compared to other AML cell lines, first implicating a functional role for *DYRK1A* in BP-MPN.^38^ Both SET- 2 and HEL have a high copy number over the *DYRK1A* locus relative to other cell lines in the Cancer Cell Line Encyclopedia (3.28 vs 1.83 respectively).^39^ HEL cells harbor a duplication of chr21q21.1-term^40^ and are a clear outlier among AML cell lines, both highly expressing *DYRK1A* and highly dependent on *DYRK1A* (CRISPR dependency score −0.72, Broad’s Cancer Dependency Map (DepMap) screening tool, **Extended Data Fig 4H**).^41–44^ Taken together, these data emphasize a role for *DYRK1A* in supporting leukemic survival, and nominate the BP-MPN cell lines HEL and SET2 as relevant models in which to study *DYRK1A*’s functional role.

## *DYRK1A* promotes proliferation and survival in chr21amp BP-MPN

We next tested the impact of *DYRK1A* knockout and knockdown using CRISPR and short hairpin (sh)RNA-approaches in HEL and SET2 BP-MPN cell lines (**Fig 5A-D**, **Extended Data Fig 5A-F**). *DYRK1A* knockout/knockdown was confirmed by Western blot (**Fig 5A and C**, **Extended Data Fig 5B and D**). *DYRK1A* targeting by both CRISPR knockout (**Fig 5B**, **Extended Data Fig 5A**) and shRNA knockdown (**Fig 5D**, **Extended Data Fig 5C and E**) significantly slowed proliferation of HEL and SET2 cell. Similarly, pharmacologic inhibition of DYRK1A using the small molecule inhibitors GNF2133 and EHT1610 led to a dose-dependent reduction in HEL cell proliferation (**Fig 5E and F**). This confirms sensitivity of JAK2 mutant BP-MPN cell lines to DYRK1A inhibition and its potential as a therapeutic target.

**Fig. 5.**
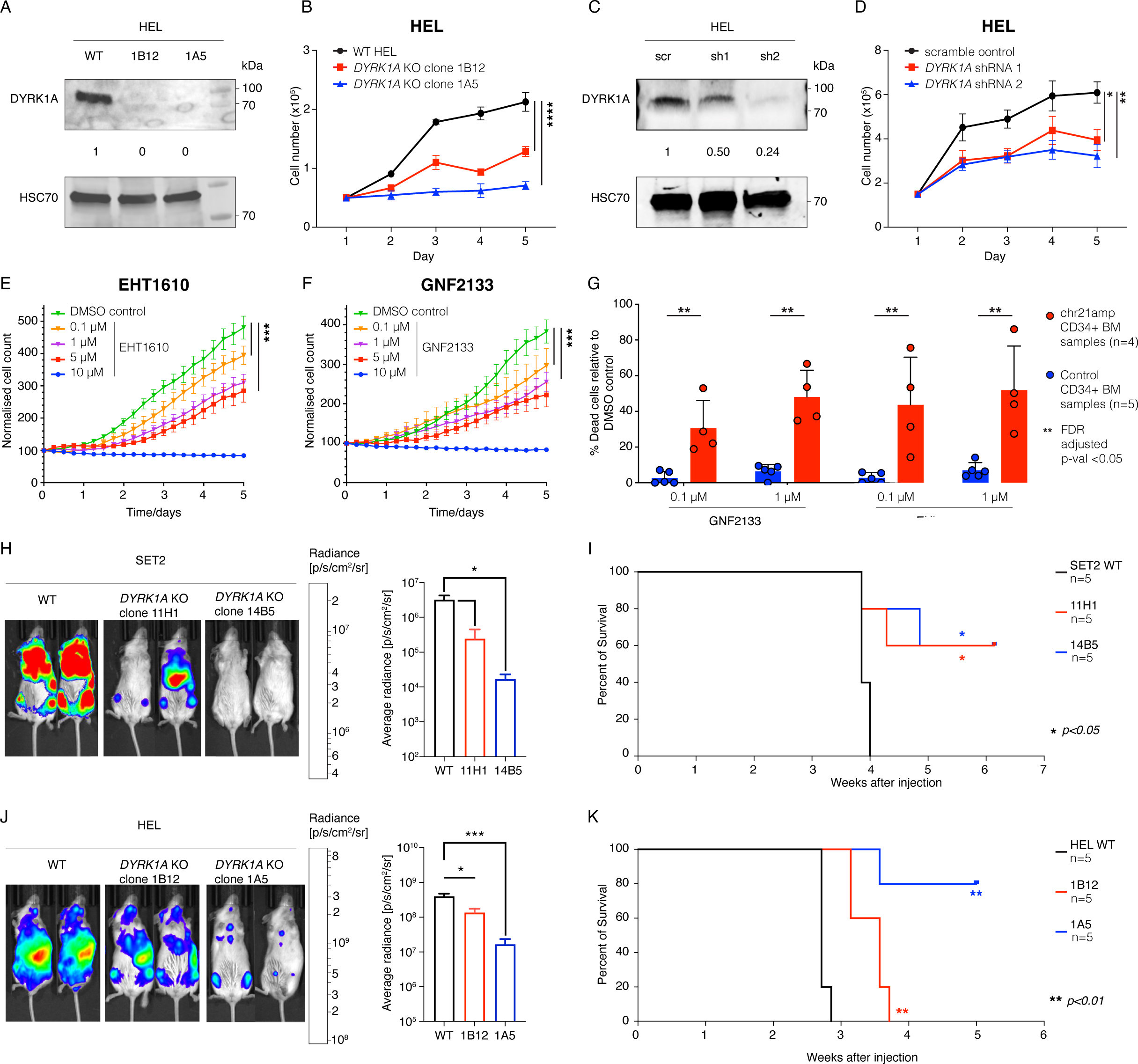
DYRK1A promotes proliferation and survival in chr21amp BP-MPN. **A.** Western blot showing the knockout of DYRK1A in HEL cells. Densitometric values were normalized to HSC70. **B.** Cell counts for HEL wild type and two *DYRK1A* KO clones (1B12 and 1A5) generated by CRISPR/Cas9 and expanded in culture (n=3 biological replicates per condition, significance calculated by 2 way ANOVA with Bonferroni’s post test) **C.** Western blot showing the knockdown of DYRK1A in HEL cells following transduction with lentiviruses expressing target specific shRNA or scramble control. Densitometric values were normalized to HSC70. **D.** Cell counts for transduced HEL cells expanded in culture (n=3 biological replicates, significance calculated by 2 way ANOVA with Bonferroni’s post test). **E. & F.** Dose-response curve treating HEL cells with increasing doses of the *DYRK1A* inhibitor EHT1610 (n=6 replicates per condition, significance calculated by 2 way ANOVA with Bonferroni’s post test) (**E**) or GNF2133 (n=6 replicates per condition, significance calculated by 2 way ANOVA with Bonferroni’s post test) **(F) G.** Primary patient BP-MPN (n=4) vs healthy control (n=5) CD34+ cell viability day 5 after treatment with 0.1 or 1 uM of GNF2133 or 0.1 or 1 uM of EHT1610 (FDR-adjusted q-value <0.05 for each). **H.** Bioluminescent images of representative mice 2 weeks following transplantation of 3 x 10e6 luciferase expressing wild-type SET2, DYRK1A KO clone 11H1 or DYRK1A KO clone 14B5 cells (n=5 each). The intensity of luminescence is normalized and shown as Avg Radiance [p/s/cm²/sr], significance calculated by unpaired t-test. **I.** Kaplan–Meier survival curves of mice (n=5 each) injected with luciferase expressing wild-type SET2, *DYRK1A* KO clone 11H1 or *DYRK1A* KO clone 14B5 cells (significance calculated by Mantel– Cox log-rank test). **J.** Bioluminescent images of representative mice 3 weeks following transplantation of 3 x 10e6 luciferase expressing wild-type HEL, *DYRK1A* KO clone 1B12 or *DYRK1A* KO clone 1A5 cells (n=5 each). The intensity of luminescence was normalized and showed as Avg Radiance [p/s/cm²/sr], significance calculated by unpaired t-test. **K.** Kaplan–Meier survival curves of mice (n=5 each) injected with luciferase expressing wild-type HEL, *DYRK1A* KO clone 1B12 or *DYRK1A* KO clone 1A5 cells (significance calculated by Mantel–Cox log-rank test). (* p<0.05; ** p<0.01; ***p<0.001; ****p < 0.0001).

To determine the relevance of these findings in primary cells, we tested the impact of DYRK1A inhibition in CD34+ HSPCs cells from patients with BP-MPN. HSPCs from 4 patients with chr21amp BP-MPN and healthy controls were treated with GNF2133 and EHT1610 at 0.1uM and 1uM doses (**Extended Data Fig 5G**). By day 5, there was a substantial and selective reduction in the viability of BP-MPN cells with DYRK1A inhibition, while cells from 5 healthy controls were unaffected (48% vs 93% viable, *padj=0.0002* for EHT1610 1uM), indicating a selective therapeutic vulnerability of chr21amp patient cells to DYRK1A inhibition (**Fig 5G** and **Extended Data Fig 5H**).

To investigate the effect of *DYRK1A* on the leukemia-propagating capacity of *JAK2* mutant BP-MPN cell lines *in vivo*, we performed CRISPR-mediated *DYRK1A* knockout in luciferase-tagged SET2 and HEL cell lines and compared their leukemogenic capacity in xenografts using immunodeficient mice (**Fig 5H-K**). The intensity of luminescence was significantly reduced in the CRISPR knockout context in both cell lines (**Fig 5H and J**), demonstrating that *DYRK1A* KO resulted in reduced ability to propagate leukemia *in vivo*. This was associated with a significant survival advantage (median survival 3.9 weeks for SET2 WT vs not reached for KO clones 14B5 and 11H1, *p=0.02* (Mantel–Cox log-rank test), median survival post-injection 2.7 weeks for HEL WT vs 3.6 weeks for KO clone 1B12 vs not reached for KO clone 1A5*, p<0.001* (Mantel–Cox log-rank test) **Fig 5I&K).**

Overall, these data validate DYRK1A as the key driver of leukemic progression in a significant proportion of patients with BP-MPN, and confirm that chr21amp confers a selective vulnerability to DYRK1A inhibition, highlighting a potential role for DYRK1A inhibitors in this context.

## *DYRK1A* contributes to the DNA damage response via regulation of the DREAM complex

BP-MPN is associated with genetic instability (**Extended Data Fig 1A**) in keeping with impaired DNA repair and/or cell survival pathways as a mechanism promoting genetic evolution. As a quarter of chr21amp cases were wild-type for *TP53*, we reasoned that the chr21amp event itself might similarly perturb DNA repair and/or cell-survival pathways.

DYRK1A-dependent phosphorylation of LIN52 is a requisite initiating step in the assembly of the DREAM complex, a key repressor of DNA repair.^45^ DYRK1A inhibitors have been shown to induce expression of DREAM-targeted DNA repair genes and promote resistance to DNA damage.^46^ Activation of the DREAM complex as a tumor-promoting and chemoresistance mechanism has been proposed in solid tumors including ovarian cancer and gastrointestinal stromal cell tumors.^47,48^

We hypothesized that *DYRK1A* overexpression in BP-MPN may enable inappropriate DREAM complex activation, with consequent transcriptional repression of DNA repair pathways. To test this, we interrogated the chr21amp and *DYRK1A* CRISPR KO cell line RNA-seq datasets for the DREAM DNA repair gene signature, derived from ChIP-seq data, identifying sites co-bound by key DREAM complex components in human cells.^46^ Indeed, *DYRK1A* CRISPR KO SET2 cells showed significant upregulation of DREAM complex target genes (**Fig 6A and B**, NES 1.76, *FWER p-value <0.001*). Moreover, in primary patient chr21amp BP-MPN cells vs controls, as well as in BEAT AML top *DYRK1A* expressors vs bottom, the DREAM DNA repair geneset was downregulated (**Fig 6C-F**, NES −1.74, *FWER p-value 0.01* for chr21amp vs non, NES −2.13, *FWER p-value <0.001* for BEAT AML top *DYRK1A* expressors vs bottom). These data support our hypothesis that DYRK1A overexpression leads to increased DREAM complex mediated transcriptional repression of DNA repair pathways in BP-MPN cells.

**Fig. 6.**
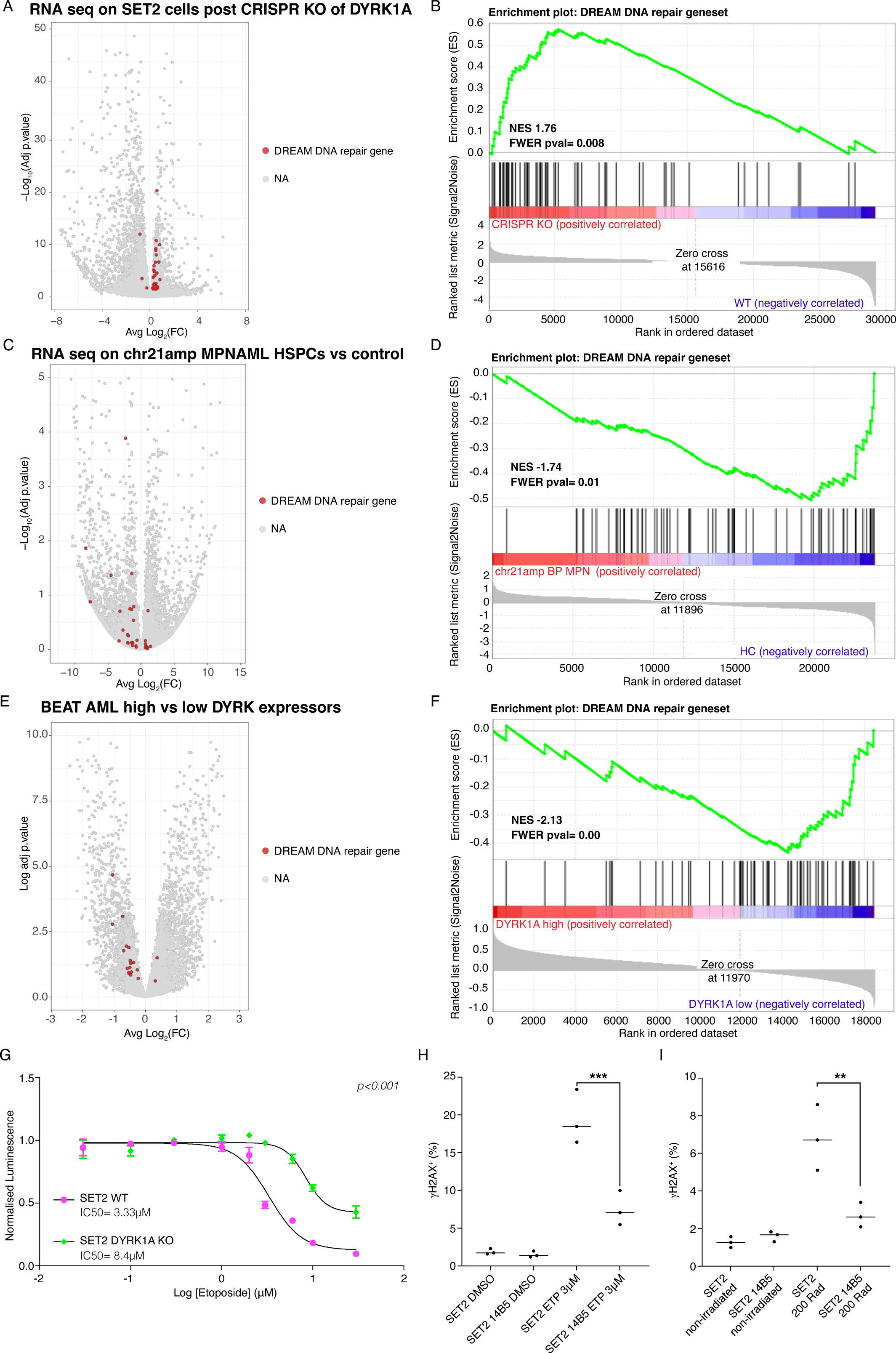
DYRK1A contributes to the DNA damage response via regulation of the DREAM complex. **A.** The transcriptional signature of DREAM complex genes involved in DNA repair is upregulated after CRISPR KO. Volcano plot of differentially expressed genes comparing CRISPR KO (n=5, 2 clones) vs WT (n=3), highlighting genes belonging to the DREAM DNA repair complex **B.** Geneset enrichment analysis demonstrating significant enrichment for DREAM DNA repair complex genes (Normalised enrichment score (NES) 1.63, Familywise-error rate (FWER) p-value 0.008). **C.** Comparing chr21amp (DYRK upregulated n=5) vs non-chr21amp (n=5) CD34+ HSPCs, the transcriptional signature of DREAM complex DNA repair genes is downregulated, which is significant on GSEA shown in **D.**(NES −1.74 FWER p-value 0.01). **E.** Comparing BEAT AML top quintile DYRK1A expressors (n=72) vs bottom quintile DYRK1A expressors (n=72), the transcriptional signature of DREAM complex DNA repair genes is downregulated, which is significant on GSEA shown in **F.**(NES −2.13 FWER p-value 0.00). **G.**Dose-response curves in CRISPR KO vs WT SET2 clones assessing proliferation by the CellTitreGlo proliferation assay after 48hrs treatment with etoposide demonstrates that loss of function of DYRK1A is protective, with higher doses required to curb cell proliferation. IC50=8.4uM for DYRK1A KO and 3.3uM for SET-2 WT clones (n=2 independent replicates, p<0.001 by Extra-Sum-of-Squares F test comparing non-linear fit curves. **H.** % cells staining positive for !- H2AX on flow cytometry at 8hrs post 3uM etoposide treatment by cell type (n=3 replicates per condition, comparison by ANOVA adjusted for multi-sample testing. *** indicates padj<0.001). **I.** % cells staining positive for !- H2AX on flow cytometry at 2hrs post 200Rad irradiation treatment by cell type (n=3 replicates per condition, comparison by ANOVA adjusted for multi-sample testing. ** padj<0.01).

To assess whether loss of *DYRK1A* in BP-MPN might restore DNA repair pathways^46^, we induced DNA damage in wild-type and *DYRK1A*-CRISPR KO SET2 cells by treatment with etoposide, a topoisomerase II inhibitor that induces double stranded DNA breaks. *DYRK1A* KO led to a shift in the dose-dependent inhibition of cell proliferation and viability, with KO cells showing greater resistance to etoposide, suggesting reduced DNA damage induced apoptosis (**Fig 6G**; *p<0.001*). This finding was supported by *DYRK1A* KO leading to a reduction in double-stranded DNA breaks as ascertained by !-H2AX staining after 8 hour treatment with 3 μM etoposide in *DYRK1A* CRISPR KO SET2 cells compared to wild type (**Fig 6H)**. Consistent with this, induction of DNA damage by irradiation in *DYRK1A* KO vs WT SET2 cells led to fewer detectable double-stranded DNA breaks at 8 hours in the KO than wild type (**Fig 6I**), which we infer may be due to enhanced kinetics of repair.

Taken together, these data support that chr21amp induced *DYRK1A* overexpression leads to suppression of DNA repair through aberrant DREAM complex activity. This might lead to increased genetic instability, in keeping with the increased number of CNAs we observed in chr21amp BP-MPN cases (**Extended Data Fig 1C**).

## *DYRK1A* upregulation is associated with activation and amplification of JAK/STAT signaling axis and the consequent upregulation of *BCL-2*

We consistently observed evidence of transcriptional upregulation of the JAK/STAT signalling axis across single cell and bulk datasets in association with chr21amp and *DYRK1A* overexpression (**Fig 4A, 4B, 4I-K, and 7A-C; Extended Data Fig 4G**). To further explore whether this might contribute to exacerbation of MPN disease phenotype, we analysed gene expression data from SET2 with and without *DYRK1A* knockout. SET2 cells showed downregulation of STAT5 target genes after *DYRK1A* CRISPR KO (NES −2.08 FWER p-value 0.001, **Fig 7A, Extended Data Fig 6A**). Furthermore, chr21amp BP-MPN cases showed enrichment of STAT3 (**Fig 7B**) and STAT5 (**Fig 7C**) gene sets in comparison with healthy controls. Consequently, we sought to investigate whether *DYRK1A* might drive disease progression through interaction with and further amplification of JAK/STAT signalling.

**Fig. 7.**
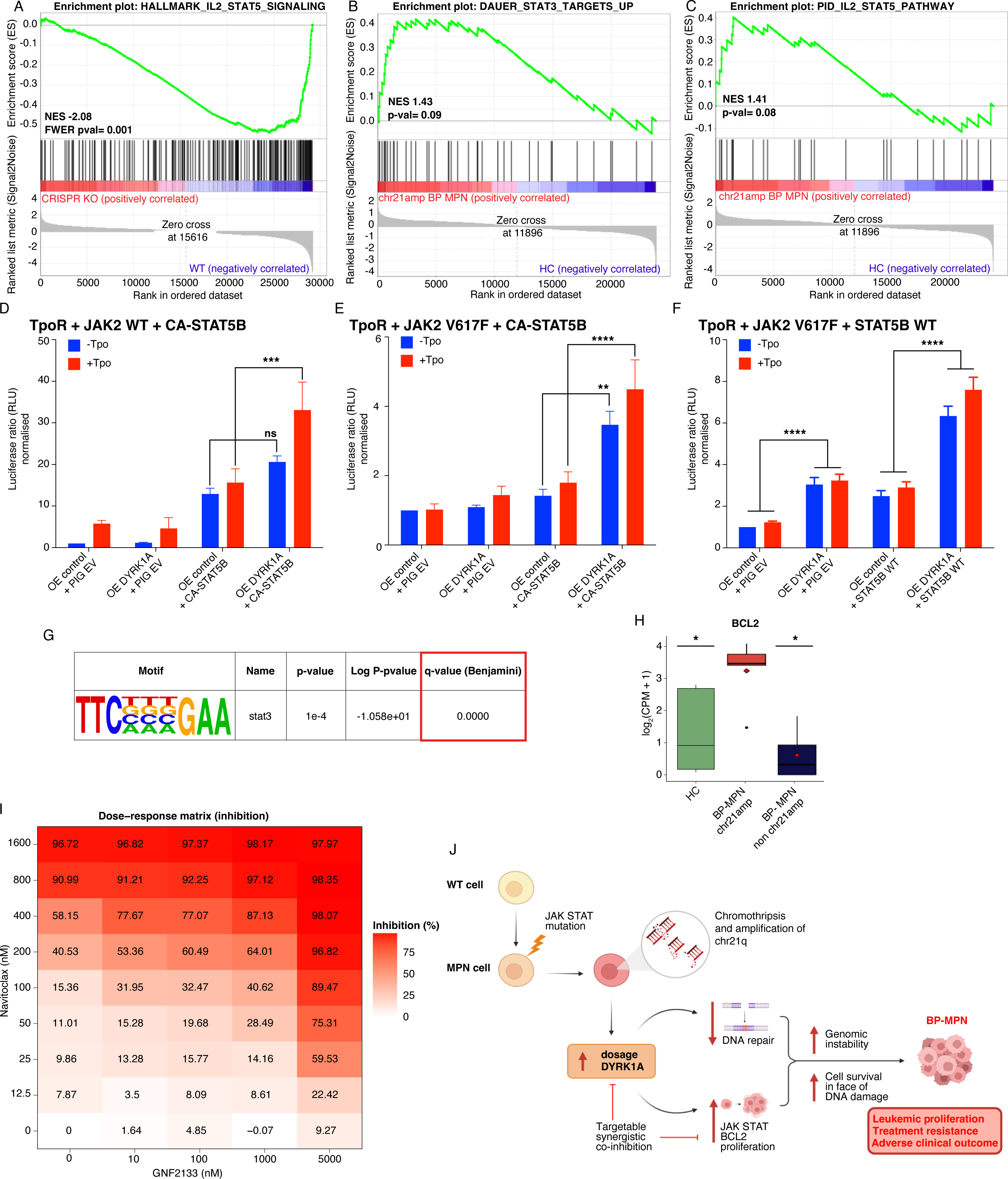
DYRK1A upregulation is associated with activation and amplification of the JAK/STAT signaling axis and the consequent upregulation of *BCL-2*. **A.** The HALLMARK IL2 STAT5 geneset is downregulated in CRISPR KO (n=5, 2 clones) vs WT (n=3) control SET2 cells, while STAT3 (**B.)** and STAT5 (**C.**) genesets are upregulated in chr21amp BP-MPN vs control cells (RNA-seq, n=5 cases per condition) **D-F** Luciferase reporter assays for STAT5 transcriptional activity in the context of *DYRK1A* wildtype overexpression versus control: HEK-293T cells were transfected with human-TpoR wild-type, murine-JAK2 wildtype and either empty vector (PIG EV) or constitutively active STAT5B (CA-STAT5) and an overexpression control (OE control) or a DYRK1A wild type overexpression vector (OE DYRK1A). At 24 hours post transfection, STAT5B dependent transcriptional activity with (red) or without (blue) TPO treatment at 6 hours was measured by the firefly luciferase assay system with Spi-Luc reporter (STAT5 response elements), as an internal control. In **E.** HEK293T cells were transfected as previous but with a mutant murine JAK2 V617F construct, and for **F.,** a wild type STAT5B vector rather than the constitutively active form was transfected. Shown are means ± SEM of three independent experiments in triplicate. Significance was assessed using Tukey’s multiple comparison test (**p<0.01, ***p<0.001,****p<0.0001, ns not significant). **G.** HOMER motif discovery analysis searching for the palindromic core STAT binding motif demonstrating significant enrichment in chr21amp peaks versus background **H.** *BCL2* expression is upregulated on RNA-seq in chr21amp BP-MPN vs controls (Paired Wilcoxon rank-sum test, \**p<0.05*). **I.** Synergy matrix scores between GNF2133 and navitoclax across a range of concentrations. Results represent mean % viability assessed by Annexin V/Propidium iodide staining by flow cytometry, normalised to DMSO-treated control wells for 6 replicates per condition. **J.** Schematic of proposed model of chr21amp driving BP-MPN transformation.

DYRK1A-dependent phosphorylation of STAT3 at Ser-727 has been shown to occur in i) an unbiased *in vitro* kinase assay seeking phosphorylation targets of DYRK1A in primary murine pre B cell lysates^49^, ii) an *in vitro* kinase screen seeking binding partners for the STAT3 Ser-727 site, which showed close correlation of the *in vitro* kinase activity of DYRK1A with *in vivo*-phosphorylation of STAT3^50^, and iii) in the Ts1Cje mouse model of Down syndrome, where increased dosage of *DYRK1A* has been linked to Ser727 phosphorylation and enhanced STAT activation.^51^ Thus STAT3 Ser-727 is a confirmed downstream target of *DYRK1A.* In line with these prior observations, we confirmed that STAT3 Ser-727 phosphorylation also occurred in the BP-MPN cell lines and showed that DYRK1A inhibition leads to a dose-dependent reduction in STAT3 phosphorylation at Ser-727 (**Extended Data Fig 6B**).

We then sought to assess whether *DYRK1A* overexpression also directly impacts STAT5B activation, using a luciferase reporter assay in which human embryonic kidney (HEK) 293T cells were co-transfected with the thrombopoietin receptor (TPO-R) and a constitutively active STAT5B (CA-STAT5B) luciferase reporter.^52^ *DYRK1A* overexpression or empty control constructs were also co-transfected alongside either *JAK2* WT or *JAK2* V617F, and cells cultured with or without TPO. As expected in this system, STAT5B transcriptional activity was activated in the presence of CA-STAT5B in the absence of TPO (**Fig 7D**). We consistently observed increased STAT5B transcriptional activity in the presence of *DYRK1A* overexpression, but not with the control vector (**Fig 7D-F**), an effect that was significantly amplified by the TPO stimulation, particularly in the context of wild-type *JAK2* (**Fig 7D**). When *DYRK1A* was overexpressed in the context of *JAK2V617F*, a mutation that results in cytokine-independent TPO-R signaling, the effect of *DYRK1A* overexpression on STAT5B transcriptional activity was further amplified and less dependent on TPO stimulation (**Fig 7E).** This was also observed with non-constitutively active STAT5B, with a 4-fold increase in STAT5 activity with *DYRK1A* overexpression (**Fig 7F**).

We then looked for evidence of transcriptional activation and STAT binding in ATAC-seq data generated from chr21amp primary patient cells. The palindromic core motif in sequences recognized by all STATs is well-described (TTCN3GAA) and was significantly enriched in chr21amp differentially accessible peaks compared to background controls (*padj <0.001*; **Fig 7G**).

A key STAT3 target is the pro-survival oncogene B-cell lymphoma 2 (*BCL2*) gene. Consistent with a functional link between *DYRK1A* and STAT transcriptional regulation, *BCL2* was one of the top 10 co-dependencies with *DYRK1A* in the DepMap database (**Extended Data Fig 6C**).^41–43,53,54^ In the *DYRK1A* CRISPR KO SET2 clones, *BCL2* was downregulated compared to control cells (log2FC −0.76, *padj < 0.001* on DeSeq2 analysis, **Extended Data Fig 6D** for log2 counts). Furthermore, in chr21amp vs non-chr21amp primary patient BP-MPN cells, we observed significant upregulation of *BCL2* RNA expression (**Fig 7H**) and chromatin accessibility (**Extended Data Fig 6E**).

The synchronized upregulation of both *BCL2* and *DYRK1A* in chr21amp cells provided a strong rationale to look for therapeutic synergy with co-inhibition of DYRK1A and BCL2. Concordantly, co-inhibition of HEL cells with the DYRK1A inhibitor GNF2133 and the BCL-2 inhibitor navitoclax demonstrated evidence of substantial therapeutic synergy (Bliss synergy score 15.02^55^, **Fig 7I/ Extended Data Fig 6F&G**).

Collectively, these data support that *DYRK1A* overexpression in the context of basal JAK/STAT activation leads to further activation and potentiation of STAT signalling, driving oncogenicity and cell survival in part by the upregulation of *BCL2*. *BCL2* can be therapeutically targeted with an inhibitor licensed for current clinical use, with clear synergy between BCL2 and DYRK1A inhibition.^55^

## Discussion

Here we describe a recurrent chromothripsis event affecting chromosome 21 in BP-MPN, uncovering a potentially actionable therapeutic vulnerability. Through integration of SNP karyotyping, whole genome sequencing, epigenetic analyses, and single cell multiomic datasets of a large BP-MPN patient cohort, together with functional studies *in vitro* and *in vivo*, we demonstrate that chr21amp occurs more frequently than other previously recognized structural variants, and leads to overexpression of *DYRK1A* which orchestrates perturbation of DNA repair, exacerbated JAK/STAT signalling and pro-survival pathways (**Fig 7J**).

It is increasingly acknowledged that CNAs are a major contributor to cancer evolution, and that patterns of aneuploidy events are non-random and tissue specific.^31,56–59^ Recent longitudinal data in Fanconi anemia patients provided a clear example that alterations in copy number, rather than point mutations, can drive clonal evolution and leukemic transformation in myeloid diseases.^60^ Across cancer there are several examples where specific aneuploidies play key carcinogenic roles, interacting with mutation profiles to drive disease phenotypes.^56,57,61^ In BP-MPN, certain CNAs are recognized as predictors of adverse outcome,^9^ but this is the first analysis to evaluate how a specific event mechanistically supports leukemic transformation. Recent data examining *TP53* mutated BP-MPN identified convergent clonal evolution with complete loss of *TP53* wild type allele acting in concert with the gain of CNAs as a requisite for transformation to BP-MPN.^19^

Intrachromosomal amplification of chromosome 21 (iAMP21) is well-described in pediatric B-cell precursor ALL (B-ALL). In this context, recurrent breakage-fusion-breakage cycles lead to the amplification of chr21q in ∼2% of B-ALL patients, and conferring high-risk disease.^62–64^ Whether iAMP21 in B-ALL and the chr21amp event we describe in BP-MPN are genetically similar or distinct remains unclear. A recent large study of 124 iAMP21 pediatric ALL cases identified that iAMP21 is an early, clonal event, comprising different patterns of chr21 amplification similar to our findings in BP-MPN, and also significantly co-occurs with mutations in the JAK/STAT signaling pathway (79/124, 64% of cases), further supporting a synergistic interaction between the two genomic events.^65,66^ We herein provide experimental evidence that this synergy likely relates to enhanced STAT signalling mediated by *DYRK1A* overexpression. We propose chr21amp as a novel biomarker of adverse clinical outcome in BP-MPN, where it synergizes with JAK/STAT pathway mutations and represents an example of “aneuploidy addiction”, akin to an oncogene addiction. This novel biological concept is supported by recent data demonstrating that genetically engineered loss of aneuploidy, in this instance generating disomic lines from trisomic 1q+ cancer cell lines, abrogates their oncogenic potential.^67^

DYRK1A’s catalytic kinase domain is conserved across eukaryotes, supporting its key biological function.^23,68,69^ *DYRK1A* has been studied in a range of disease contexts, most closely in Down syndrome (DS) where congenital *DYRK1A* upregulation occurs by trisomy and has been implicated in the cardiac, hematological and neurological features of DS.^23,70–72^ Further diverse roles have been proposed for DYRK1A, including the phosphorylation of tau and amyloid proteins and in regulating pancreatic beta cell proliferation, leading to its investigation as a modulator of diseases ranging from Alzheimer’s to diabetes.^71,73–75^ Consequently, several DYRK1A inhibitors have been developed which show on-target efficacy and negligible toxicity in murine models and in a pilot human study, rendering it a key target for further development.^76–78^ Inhibition of DYRK1A has also recently been reported to alter splicing and increase sensitivity to BCL2 inhibition in non-DYRK1A amplified AML.^79^

Here, we propose that two biological pathways activated by *DYRK1A* are crucial in its oncogenic activity when upregulated in chr21amp BP-MPN. First, we suggest that the chr21amp event causes downregulation of DNA repair pathways to promote genomic instability. This is in accordance with a molecular events in Fanconi Anemia, where it is established that congenital loss of the Fanconi DNA repair pathway followed by *TP53* downregulation drives a survival advantage and stem/progenitor cell survival in leukemia development.^60,80^ Notably, these events occur in a different order to our observations in BP-MPN, where *TP53* mutation precedes *DYRK1A* mediated repression of DNA repair pathways.

Second, we demonstrate that *DYRK1A* upregulation leads to exacerbation of JAK2V617F-driven upregulation of JAK/STAT signalling. Our chr21amp BP-MPN cohort shows clear parallels to the B-cell ALL iAMP21 clinical subgroup, which is characterized by an enrichment in JAK/STAT signalling and shares the same region of amplification.^63–66^ Our interrogation of published *de novo* AML datasets for the chr21amp event confirms its co-occurrence with *TP53* and rarity outwith the JAK/STAT mutant setting, corroborating a small case series of chr21amp AML patients.^81^ Taking our functional analyses together with the observation from CRISPR screening that AML lines harboring *JAK2* mutations are hypersensitive to targeting of *DYRK1A* ^38,43^, we conclude that in chr21amp BP-MPN there is synergy and potentiation between *DYRK1A* overexpression and basal JAK/STAT upregulation. This further potentiation of an oncogenic pathway already activated by mutation by CNAs is documented in other cancer settings, such as in the case of BRAF mutant solid tumors where acquisition of CNAs lead to further aberrant amplification of the MAP kinase pathway.^82^

In summary, we describe a high frequency of chromothripsis and other structural variants in BP-MPN, and identify a recurrent region of amplification of chromosome 21 as a novel prognostic biomarker. Through multiomic analysis of patient samples coupled with *in vitro* and *in vivo* functional assays, we describe how chr21amp creates a therapeutic vulnerability in BP-MPN through a druggable DYRK1A-BCL2 axis. This provides a paradigm for the translation of recurrent regions of aneuploidy to an actionable molecular target.

## Acknowledgements

We are grateful to patients and donors, without whom this study would not have been possible. We thank Patricia Ciccone, Nawshad Hayder and Nikos Sousos who helped with sample banking; Kevin Clark, Craig Waugh and Paul Sopp in the MRC WIMM Flow Cytometry facility which is supported by the MRC Human Immunology Unit and MRC Molecular Haematology Unit; Dr. Neil Ashley in the MRC WIMM Single Cell Facility; Ryan Beveridge in the MRC WIMM Virus Screening Facility, Lance Palmer at St Jude Children’s Research Hospital, and all the staff at Isabl Inc for facilitating the whole genome sequencing analyses. We also thank the Oxford Genomics Centre at the Wellcome Centre for Human Genetics (funded by Wellcome Trust grant reference 203141/Z/16/Z) for the generation and initial processing of the OmniExpress SNP array data. This work was supported by a CRUK Advanced Clinician Scientist Fellowship (to B.P, Grant number C67633/A29034), a CRUK Senior Cancer Research Fellowship (to A.J.M., Grant number C42639/A26988), and a Sir Henry Wellcome Clinical Doctoral Training Fellowship (to CKB; 220586/Z/20/Z). The authors would like to acknowledge the National Institute for Health Research (NIHR), Oxford Biomedical Research Centre (BRC); John Fell Fund (131/030 and 101/517), the EPA fund (CF182 and CF170) and by the MRC WIMM Strategic Alliance awards G0902418 and MC_UU_12025, and the contribution of the WIMM Sequencing Facility, supported by the MRC Human Immunology Unit and by the EPA fund (CF268). Additional funding was provided by the National Institutes of Health (NIH) grants (numbers R35CA253096 and P01CA108671), the MPN Research Foundation (to JDC and AT), the Samuel Waxman Cancer Research Foundation (to JDC), Syndax Pharmaceuticals (to JDC), and St. Jude/ALSAC (to JDC). The views expressed are those of the authors and not necessarily those of the National Health Service (NHS), the NIH, the NIHR or the Department of Health. In addition, this research was funded by Cancer Research UK RadNet Manchester (C1994/A28701) and supported by the NIHR Manchester Biomedical Research Centre (NIHR203308). The results published here are in part based upon data generated by the TCGA Research 1250 Network (https://www.cancer.gov/tcga) and the BeatAML team. The views expressed are those of the author(s) and not necessarily those of the NIHR or the Department of Health and Social Care.

---

**Extended Data Fig 1.**
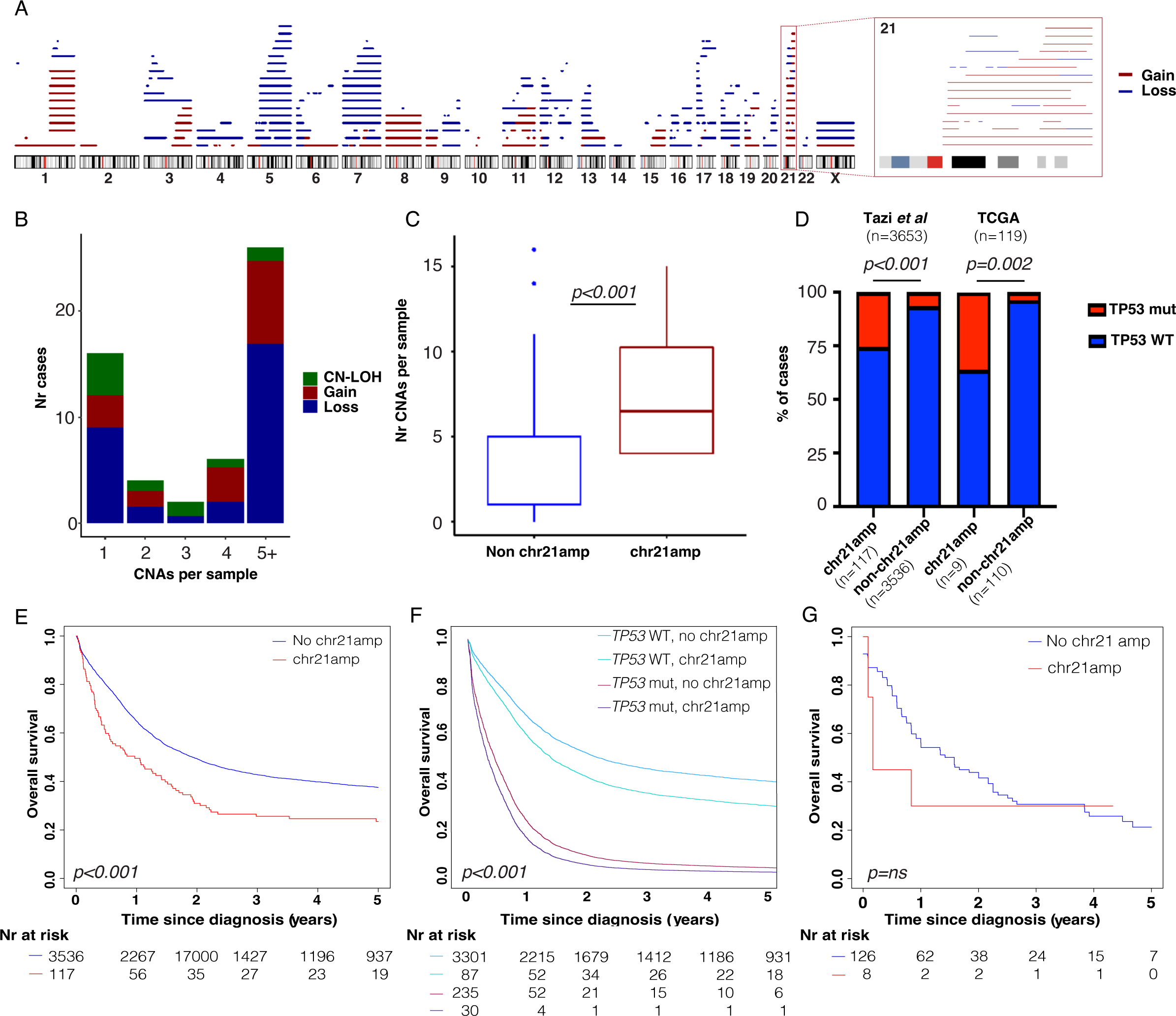
Supplemental information on copy number alterations in BP-MPN cohort and chr21amp in external AML datasets. **A.** Graphical representation of all copy number gains and losses identified across the cohort by MoChA.^25,26^ Events are colored by type (gain=red, loss=blue) and overlay chromosomal ideograms to indicate location. **B.** Number of cases by number of chromosomal alterations. Events are colored by event type (gain=red, loss=blue, CN-LOH=green). **C.** chr21amp cases have a greater number of non-chr21 copy number abnormalities compared to non-chr21amp cases (median= 6.5(IQR 4-10.3) vs median 1(IQR 1-5), *p<0.001* by Wilcoxon rank-sum test). **D.** In both the TCGA L-AML cohort and a large cohort of 3653 *de novo* AML patients, there was a significant enrichment for mutated and/or deleted *TP53* amongst chr21amp cases (Fisher’s exact test, *p<0.001* for both) **E.** Kaplan Meier survival curve stratified by chr21amp status showing significantly impaired overall survival (OS) for chr21amp amongst 3653 *de novo* AML cases (median OS from diagnosis 0.97 (95%CI 0.55-1.47) vs 1.93years (95% CI 1.77-2.09) *p<0.01* by Mantel Cox log-rang test). **F.** Cox survival curve stratifying patients in (b) by *TP53* and *chr21amp* status, showing that the adverse impact on survival conferred by chr21amp persists after stratifying by *TP53* status (*p=0.009*, HR for chr21amp alone 1.3 (95% CI 1.1-1.7), mutant TP53 alone 3.75 (95%CI 3.3-4.3), HR for mTP53 with chr21amp= 4.71 (95%CI 3.3- 6.8) **G.** The TCGA dataset is underpowered for a survival analysis by chr21amp (median OS in years chr21amp 0.17 years (95%CI 0.17-NA) vs no chr21amp 1.58 years (95%CI 0.92-2.2), *p=0.4)*

**Extended Data Fig 2.**
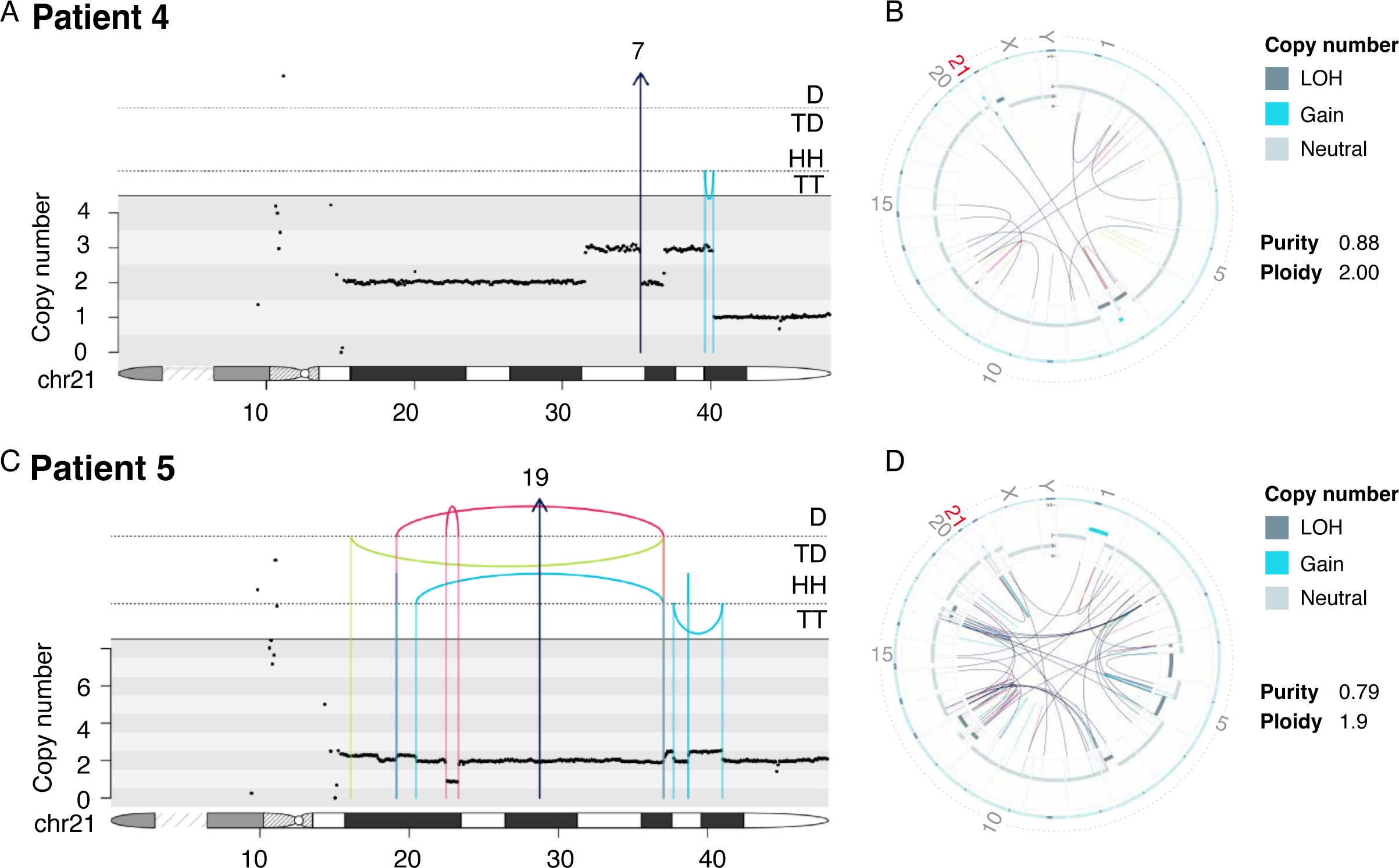
Whole genome sequencing extended figures. **A.& C.** integrated CN & structural variant plots for patients 4 & 5. As in Fig 2, the top panel shows intra-chromosomal events as arcs between breakpoint loci, and color denotes the type of SV (black=translocation, red=deletion, blue=duplication, green=inversion). Rearrangements are further separated and annotated based on orientation. D: Deletion, TD: Tandem duplication; HH, head-to-head inverted, TT: tail-to-tail inverted. Inter-chromosomal events are shown with arrows denoting the likely partner chromosome. The middle panel shows the consensus copy number across the chr21 ideogram, depicted in the lowest section of each plot to indicate breakpoint location. Patient 4 has a simple breakage-fusion-bridge event, whereas patient 5 shows a low-level copy number gain and inversion event. **B.&D.** Circos plots for patients 4 & 5 showing global SV burden, demonstrating clustering around chr21. The outer ring shows the chromosome ideogram. The middle ring shows the B allelic frequency and the inner ring shows the intra-and inter-chromosomal SVs with the same color scheme as in **A&C**.

**Extended Data Fig 3.**
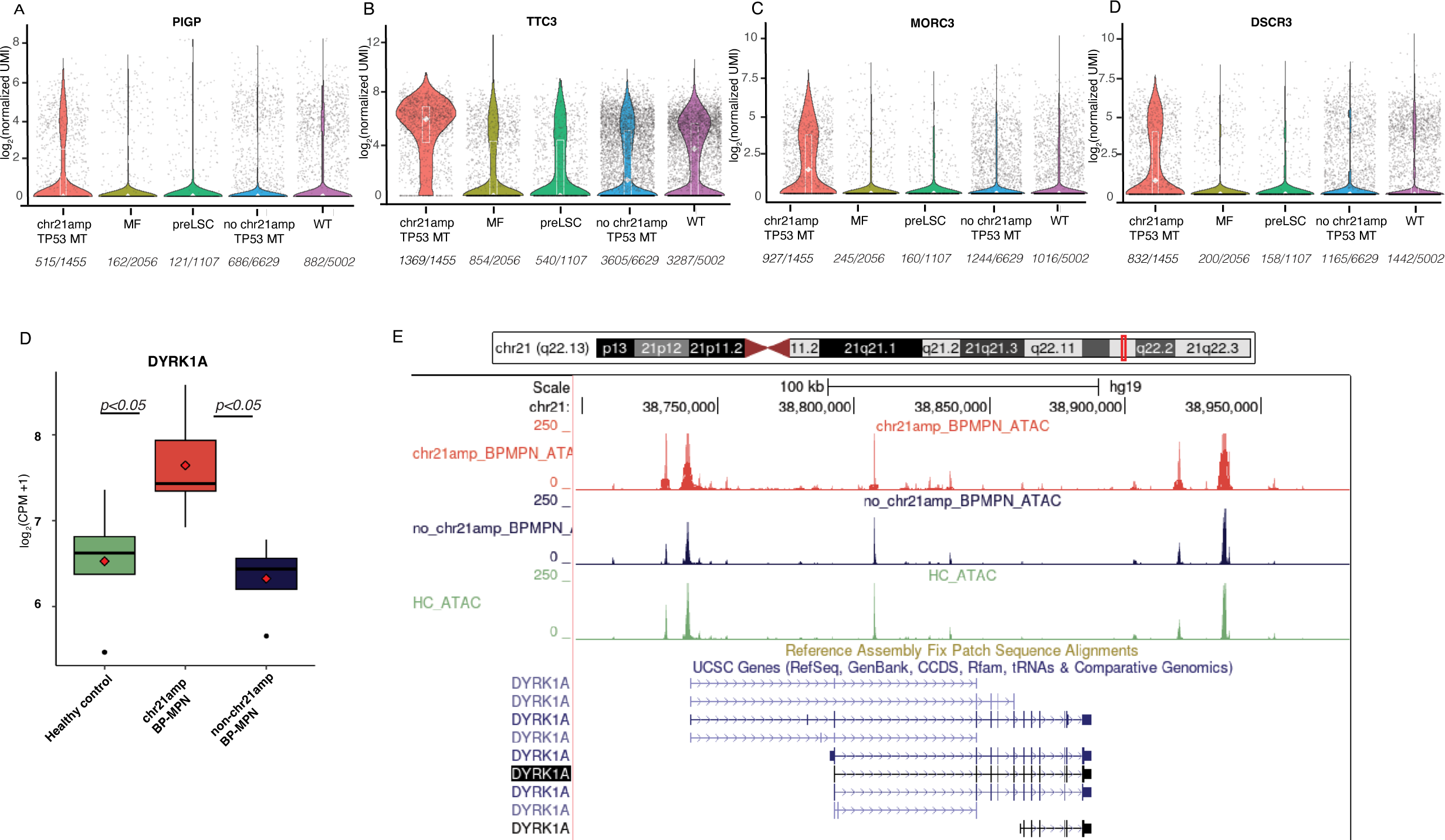
Supplemental information on prioritising genes in the minimally amplified region using TARGET-seq. **A.-D.** Violin plots of the significantly DE genes (log2FC>1, padj<0.05) in chr21amp HSPCs compared to non-chr21amp control cells including myelofibrosis (MF, n-2056 cells from 8 MF donors), pre-leukemic stem cells (n=1107 non-mutant phenotypic HSC, identified in 12 BP-MPN donors), non-TP53-non-chr21amp BP-MPN (n=6629 cells from 14 donors) and wild type cells (n=5002 from 9 healthy donors). Each dot represents the expression value (log2-normalized UMI count) for each single cell, with median and quartiles shown in white. Expressing cell frequencies are shown on the bottom of each violin plot for each group **A.** PIGP **B.** DSCR3 **C.** MORC3 **D.** TTC3. For each gene, *padj<0.001* when comparing chr21amp_TP53_MT vs all other categories by Wilcoxon rank sum test, adjusted for multiple comparisons by Bonferroni correction. **E.** Plot of normalised counts (log2CPM+1) of DYRK1A expression on RNA_seq analysis, showing mean (diamond), median and IQR. *DYRK1A* is overexpressed in chr21amp cases compared to controls (Wilcoxon rank-sum test, *p<0.05* for all comparisons). **F.** ATAC-seq tracks shown in UCSC genome browser window over the *DYRK1A* genomic location by condition (chr21amp in red, healthy control in green, non-chr21amp BP-MPN in blue).

**Extended Data Fig 4.**
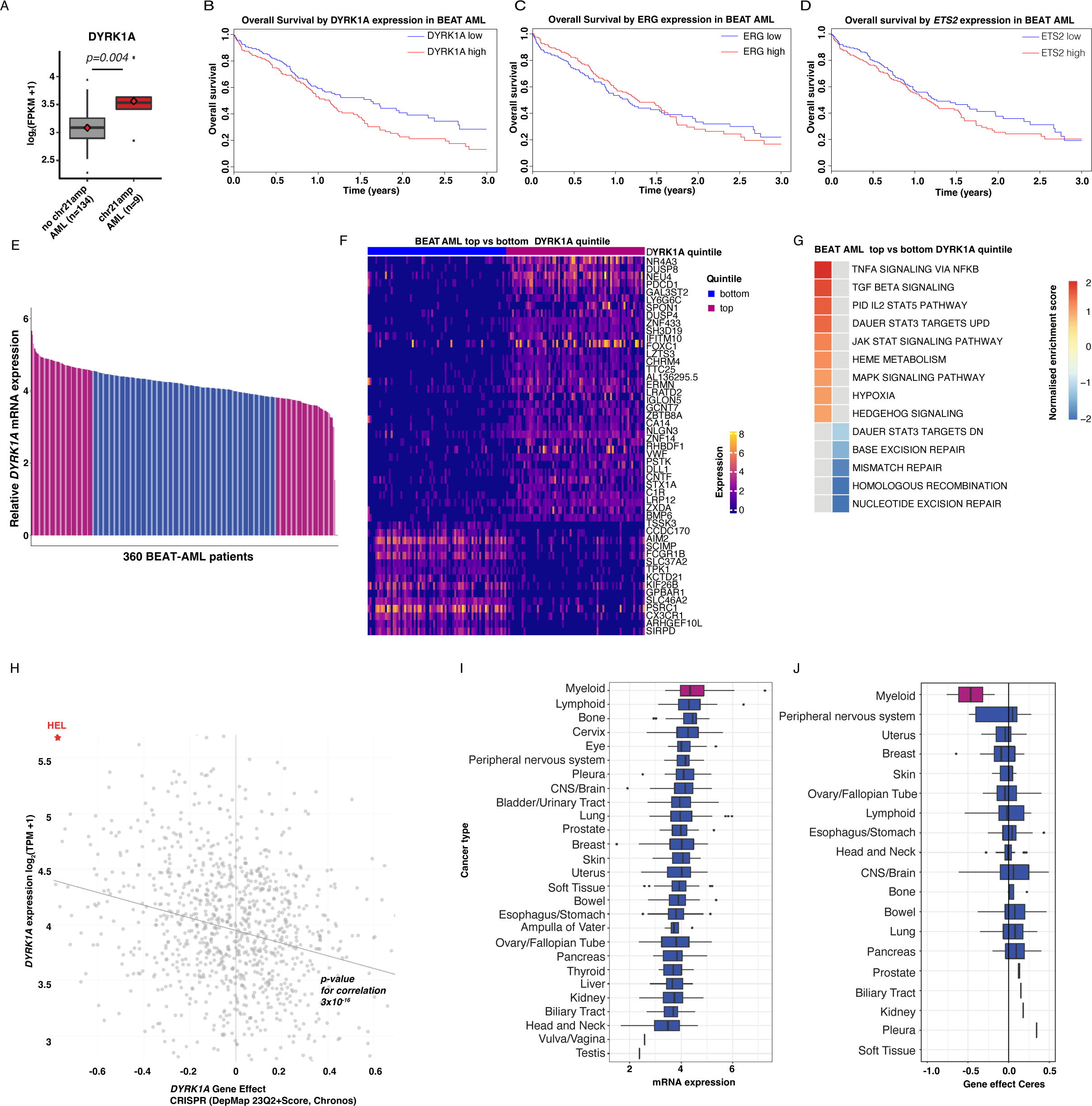
*DYRK1A* expression in external AML datasets. **A**. Gain of chr21q in the TCGA cohort leads to upregulation of *DYRK1A* expression (median log2FPKM 3.53 vs 3.08 [IQR 3.32-3.74 vs 2.73-3.44], *p=0.004)* **B.** In the 360-patient BEAT AML cohort, overexpression of *DYRK1A* is associated with adverse outcome (HR 1.44 (95% CI 1.07-1.93, *p-value 0.03), Cox regression analysis,* 3year (y) OS 13.1% (95%CI 7.1-24.4%) vs 28.4%(95%CI 18.8-43.1%, *p=0.02*, Mantel–Cox log-rank test), while other genes in the chr21amp minimally amplified region are not (ERG 3y OS ERG high 16.7% (95%CU 9.3-30.1%) vs 22.0% low (95%CI 13.7-35.4%), *p=0.8* Mantel–Cox log-rank test) **C,** ETS2 3y OS ETS high 19.1%(95%CI 13.2-31.2%) vs 20.3% ETS low (95%CI 10.5-34.8%, *p=0.2* Mantel–Cox log-rank test) **D.) E.** Graphic of stratifying the BEATAML cohort by top (n=72) vs bottom (n=72) quintile of *DYRK1A* expression **F.** Heatmap of top differentially expressed genes of top vs bottom quintiles of the BEAT AML cohort stratified by *DYRK1A* expression. **G.** Hallmark and KEGG pathway GSEA of top altered pathways (NES, Normalized enrichment score >/<1) comparing patients in (**E.) H.** Plot of *DYRK1A* gene expression vs gene dependency from the Broad DepMap database, showing the chr21amp MPNAML cell line HEL as an outlier. Linear regression analysis shows correlation between expression and dependency (Pearson correlation coefficient −0.242, slope −5.67E-1, *p-value <0.001)* **I.** mRNA expression of *DYRK1A* by cell line lineage in the DepMap database **J.** Ceres gene dependency scores for *DYRK1A* by cell line lineage in DepMap

**Extended Data Fig 5.**
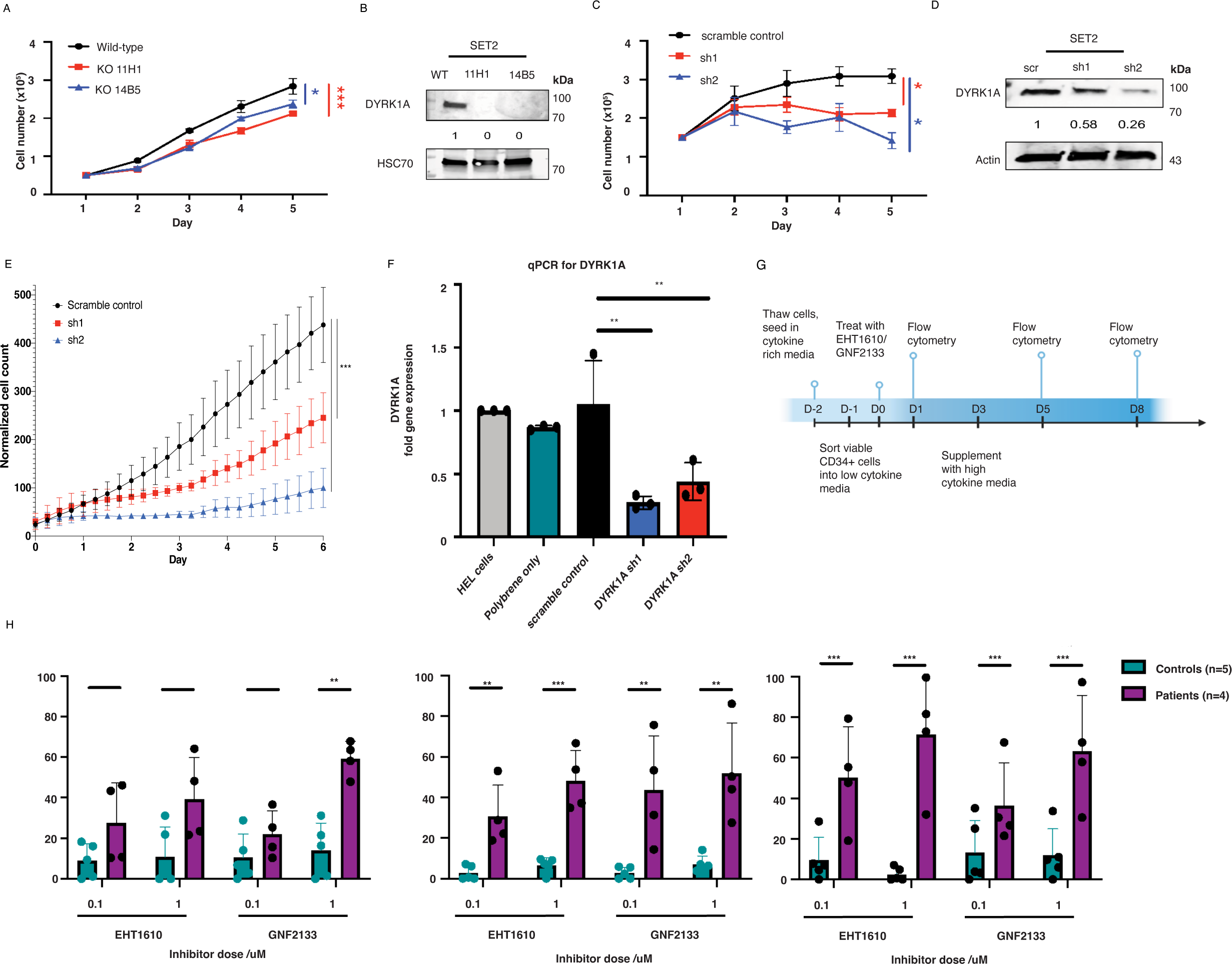
Supplemental experiments functionally validating *DYRK1A* as a target in BP-MPN. **A.** Cell counts for two SET2 *DYRK1A* KO clones (14B5 & 11H1) generated by CRISPR/Cas9 and expanded in culture, confirming adverse effect on proliferation by *DYRK1A* KO over time (n=3 biological replicates, *p<0.05* by day 5). **B.** Western blot showing the knockout of *DYRK1A* in the 14B5 & 11H1 SET2 cell clones. Densitometric values normalized to HSC70. **C.** Cell proliferation assay for SET2 cells transduced with lentiviruses expressing *DYRK1A* specific shRNA or scramble control (n=3 biological replicates, *p<0.05* by day 5). **D.** Western blot showing the knockdown of *DYRK1A* expression in SET2 cells following transduction with lentiviruses expressing target specific shRNA or scramble control. Densitometric values were normalized to actin. **E.** Independent replicate experiment of shRNA knockdown experiments. HEL cells were treated with lentiviruses expressing *DYRK1A* specific shRNA or scramble control, and placed in the Incucyte for serial cell counts (summary of 3 biological replicates, *p<0.05*). **F.** Knockdown of *DYRK1A* was validated by qRT-PCR. **G.** Experimental layout for primary patient experiments. **H.** Timecourse of viability readouts for primary patient cells on days 1,5,8 post treatment with DYRK1A inhibitors EHT1610 and GNF2133 at 0.1 and 1uM doses. On the y axis is % cell death relative to DMSO control. Each sample n=5 controls, n=4 patients) is shown by a dot, with mean and SD depicted in the boxplot. Significance testing by t-test with Bonferroni correction for multiple testing, shown are the adjusted q values **<0.01, ***<0.001.

**Extended Data Fig 6.**
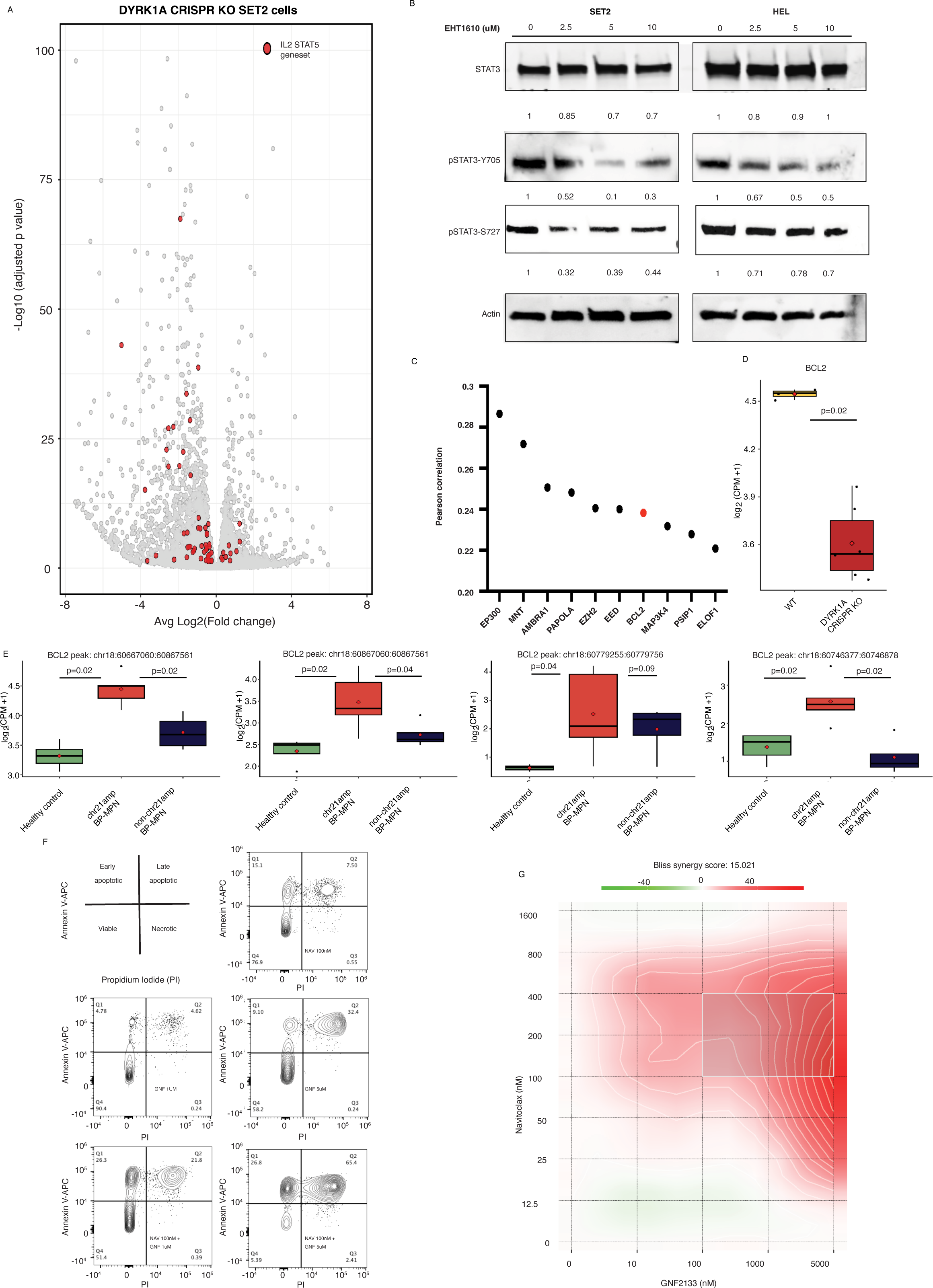
Supplemental experiments evaluating the mechanism of *DYRK1A* and the concomitant upregulation of BCL2, supporting a rationale for therapeutic synergy with co-inhibition of DYRK1A and BCL2. **A.** Volcano plot of differentially expressed genes on RNA-seq of SET2 cell lines who underwent CRISPR KO of *DYRK1A* (n=5, 2 clones) vs WT (n=3) highlighting downregulation of HALLMARK STAT5 signalling pathway genes (GSEA NES −2.08, FWER p-value <0.01) **B.** Western blot for phosphorylation of STAT3 at Y705, S727 and total STAT3 with actin normalization, at baseline and after 30 minutes of treatment with increasing doses of the DYRK1A inhibitor EHT1610 in HEL and SET2 cells **C.** Co-dependencies for *DYRK1A* in DepMap, highlighting *BCL2* amongst the top 10 genes co-dependent on CRISPR screen (Pearson’s correlation co-efficient 0.24) **D.** DYRK1A KO SET-2 cell line clones downregulate *BCL2 (*n=5 CRISPR KO vs n=3 WT*, p-0.02 Wilcoxon rank-sum test)* **E.** ATAC-seq peaks in chr21amp vs non-chr21amp cells showing upregulation of peaks across the *BCL2* gene body in chr21amp patients (n=5 chr21amp vs n=4 non-chr21amp vs n=5 healthy controls, *paired Wilcoxon rank-sum test*) **F.** Representative flow cytometry plots for Annexin V PI apoptosis assay demonstrating synergy between the DYRK1A inhibitor GNF2133 (‘GNF”) and the bcl-2/bcl-xL inhibitor navitoclax (“NAV”). The middle left panel shows that GNF1 uM induces apoptosis in 9.4% of cells while NAV 100nM does so in 22.6% of cells (top right). In combination (bottom left), NAV 100+GNF induce apoptosis in 46.1%. **G.** Bliss synergy score and matrix contour plot highlighting areas of greatest synergy between DYRK1A and navitoclax inhibitor dosing.

**Extended Data Fig 7.**
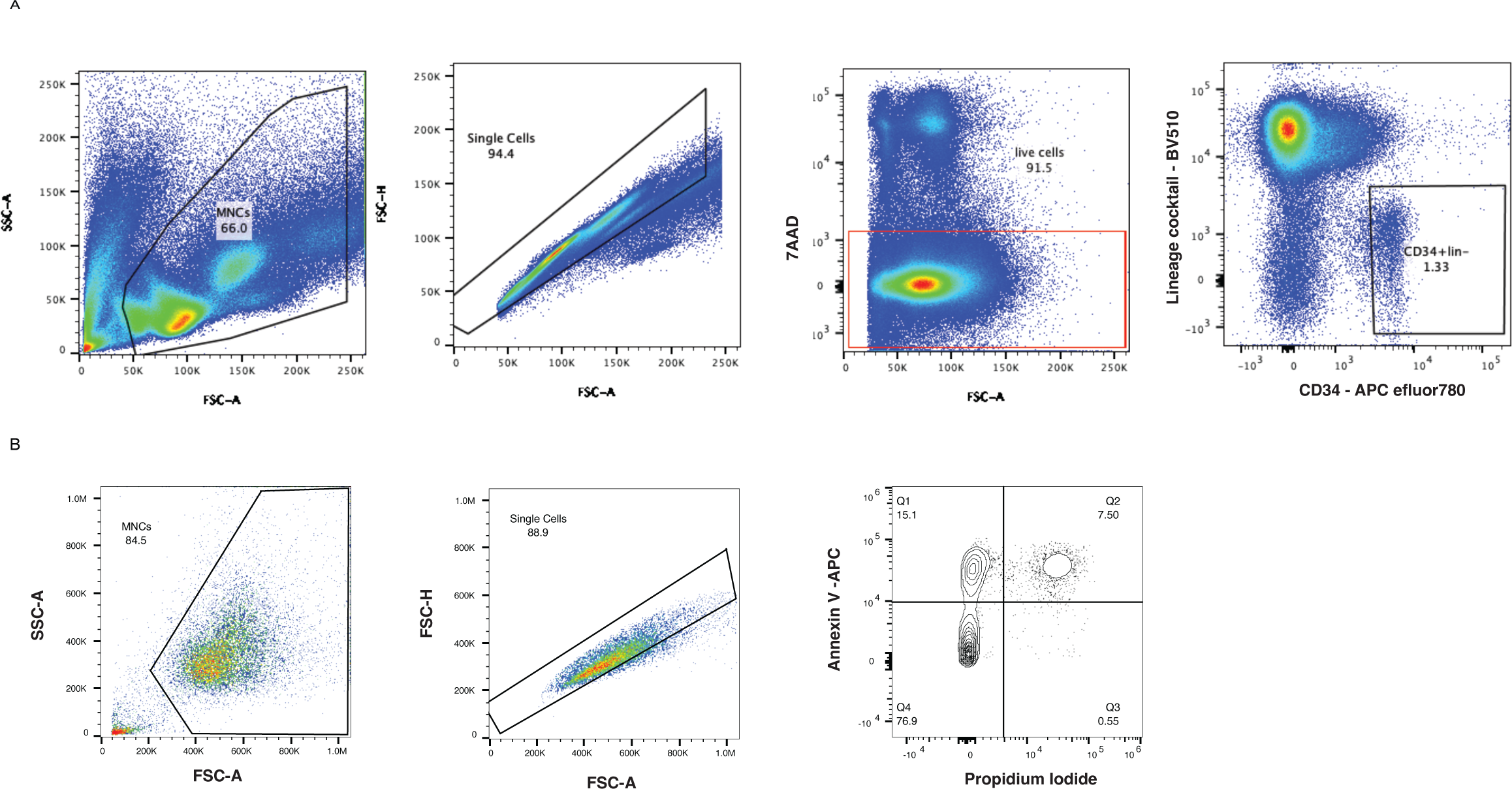
**A.** Sorting strategy for mini-bulk RNA, ATAC and 10X single cell RNA-seq experiments: Lineage^-^CD34^+^ cells were sorted Eppendorf tubes for subsequent library preparation. **B.** Gating strategy for Annexin V/PI apoptosis staining experiments. MNCs: Mononuclear cells, FSC-A: Forward Scatter Area, SSC-A: Side Scatter Area, FSC-H: Forward-Scatter Height, 7-AAD: 7- aminoactinomycin D.

## Author Contributions

**C.K.Brierley:** conceptualization,investigation, validation, funding acquisition, methodology, computational analyses, writing–original draft and editing. **O. Yip:** investigation, validation, visualization, writing–editing. **G. Orlando:** investigation, validation, writing–editing. **H. Goyal:** investigation. **W. Wen**: supervision and implementation of computational analysis. **J. Wen**: investigation, validation, visualization, writing–editing. MF Levine: implementation of computational analysis. **A. Rodriguez-Meira**: investigation **A. Adamo**: Investigation **M. Bashton**: implementation of computational analysis**. A. Hamblin**: resources, investigation. **SA Clark**: investigation, methodology. **J. O’Sullivan**: resources. **L Murphy**: resources. **A Olijnik**: writing-figure editing. **A Cotton**: investigation. **S. Pruett-Miller**: resources. **S. Narina**: resources. **A. Enshaei**: investigation. **C. Harrison**: resources. **M. Drummond**: resources. **S. Knapper**: resources. **A.Tefferi:** conceptualization. **I Antony-Debré**: resources. **S. Thongjuea**: software. **S. Constantinescu**: methodology, investigation. **E. Papaemmanuil**: Supervision, resources, investigation. **B. Psaila**: Conceptualization, resources, formal analysis, supervision, funding acquisition, methodology, writing–original draft and editing, project administration. **JD. Crispino**: Conceptualization, resources, formal analysis, supervision, funding acquisition, methodology, writing–editing. **AJ. Mead**: Conceptualization, resources, formal analysis, supervision, funding acquisition, methodology, writing–original draft, project administration. All authors read and approved the submitted manuscript.

## Inclusion and Diversity statement

We support inclusive, diverse, and equitable conduct of research.

